# Chemokine positioning determines mutually exclusive roles for their receptors in extravasation of pathogenic human T cells

**DOI:** 10.1101/2023.01.25.525561

**Authors:** Farhat Parween, Satya P. Singh, Hongwei H Zhang, Nausheen Kathuria, Francisco A. Otaizo-Carrasquero, Amirhossein Shamsaddini, Paul J. Gardina, Sundar Ganesan, Juraj Kabat, Hernan A. Lorenzi, Timothy G. Myers, Joshua M. Farber

**Affiliations:** Laboratory of Molecular Immunology, National Institute of Allergy and Infectious Diseases, National Institutes of Health, Bethesda MD 20892, USA; Research Technologies Branch, Genomic Technologies, National Institute of Allergy and Infectious Diseases, National Institutes of Health, Bethesda MD 20892, USA; Research Technologies Branch, Biological Imaging, National Institute of Allergy and Infectious Diseases, National Institutes of Health, Bethesda MD 20892, USA; Bioinformatics and Computational Biosciences Branch, National Institute of Allergy and Infectious Diseases, National Institutes of Health, Bethesda MD 20892, USA

## Abstract

Pro-inflammatory T cells co-express multiple chemokine receptors, but the distinct functions of individual receptors on these cells are largely unknown. Human Th17 cells uniformly express the chemokine receptor CCR6, and we discovered that the subgroup of CD4^+^CCR6^+^ cells that co-express CCR2 possess a pathogenic Th17 signature, can produce inflammatory cytokines independent of TCR activation, and are unusually efficient at transendothelial migration (TEM). The ligand for CCR6, CCL20, was capable of binding to activated endothelial cells (ECs) and inducing firm arrest of CCR6^+^CCR2^+^ cells under conditions of flow - but CCR6 could not mediate TEM. By contrast, CCL2 and other ligands for CCR2, despite being secreted from both luminal and basal sides of ECs, failed to bind to the EC surfaces - and CCR2 could not mediate arrest. Nonetheless, CCR2 was required for TEM. To understand if CCR2’s inability to mediate arrest was due solely to an absence of EC-bound ligands, we generated a CCL2-CXCL9 chimeric chemokine that could bind to the EC surface. Although display of CCL2 on the ECs did indeed lead to CCR2-mediated arrest of CCR6^+^CCR2^+^ cells, activating CCR2 with surface-bound CCL2 blocked TEM. We conclude that mediating arrest and TEM are mutually exclusive activities of chemokine receptors and/or their ligands that depend, respectively, on chemokines that bind to the EC luminal surfaces versus non-binding chemokines that form transendothelial gradients under conditions of flow. Our findings provide fundamental insights into mechanisms of lymphocyte extravasation and may lead to novel strategies to block or enhance their migration into tissue.

## Introduction

Effector/memory T cells provide protection against infection and mediate damage in peripheral tissue and as such are critical components of both host defense and immune-mediated disease ^1–5^. Effector/memory CD4^+^ T cells provide help for B cells, CD8^+^ T cells and mononuclear phagocytes and have irreplaceable roles in host defense in humans ^6, 7^. The resting memory Th population consists both of cells that are resident (Trm) providing more immediate immune protection ^4, 8–10^ and circulating memory T cells, which migrate from blood and peripheral tissue and rapidly respond to the invading pathogens ^11, 12^. These memory cells reflect the complexity of the cells produced during the acute response, serve as a record of effector cell differentiation, and contain cells capable of exercising immediate effector function. They can be categorized according to multiple overlapping features, such as the ability to provide B cell help in germinal centers, pathways of migration under homeostatic conditions, or patterns of cytokine production ^6^. CD4^+^ effector/memory T cells can differentiate into subsets characterized by their cytokine profiles and master transcription factors ^13, 14^. The master transcription factor for Th1 cells is T-bet, the signature cytokine is IFN-γ, and these cells offer protection primarily against intracellular bacteria and viruses, whereas for Th17 cells the master transcription factor is retinoid-related orphan receptor gamma-t (RORγt), the signature cytokine is IL-17A, and these cells protect mucosal surfaces from extracellular pathogens and mediate tissue damage in autoimmune reactions ^13, 14^. Although “Th17” refers to Th cells that can produce IL-17A/F, the Th17 family of cells can be viewed more broadly to include, for example, some cells expressing RORγt that can produce IL-22, CCL20 and IFN-γ, with or without IL-17A/F, and some of these cells have been characterized as pathogenic in mouse models of autoimmune disease through their production of IFN-γ and/or GM-CSF ^15, 16^. An integral component of Th cell pathogenicity is the ability to traffic into and within tissue, and this capability depends on the activities of chemoattractant receptors. One such receptor, CCR6, is expressed on all human T cells that can produce IL-17A/F and the other proteins associated with the Th17 family, and we have used CCR6 to identify this greater population that we include in the category of “type 17” cells ^17^.

Consistent with its pattern of expression and activity, CCR6 has been shown to be important in mouse models of autoimmune disease such as experimental autoimmune encephalomyelitis and psoriasis-like inflammation of the skin ^18, 19^. Whereas most chemokine receptors bind multiple chemokines and vice versa, CCL20 is the only chemokine ligand for CCR6 and CCR6 is the only known receptor for CCL20 ^20, 21^. Despite this lack of receptor:ligand promiscuity, redundancy in the activity of trafficking receptors can be important on CCR6- expressing Th cells, since these cells typically express multiple chemokine receptors such as CCR2, CCR4, CCR5, CXCR3 and CXCR4 in various combinations (^22^ and this manuscript). This redundancy may account for examples of CCR6-independent positioning of Th17 cells at epithelia ^23, 24^ and chemokine receptor redundancy is often invoked as an explanation for disappointing results in trials of receptor antagonists in inflammatory diseases^25^.

Extravasation of leukocytes across inflamed endothelium can be separated into steps of rolling mediated by selectins, firm arrest resulting from chemoattractant receptor-induced integrin activation, and chemokine-dependent transendothelial migration (TEM) ^26^. In our previous work with human MAIT cells, we found that CCR6 could mediate firm arrest while CCR2 was responsible for TEM ^27^. In the current work we sought to investigate the activities of these and other receptors on human CCR6^+^/type 17 Th cells. We found that CCR2 identified a subgroup of CCR6^+^ cells enriched for a pathogenic pattern of cytokine production, and that these CCR6^+^CCR2^+^ T cells could produce IFN-γ and GM-CSF after culturing with cytokines alone, without activation through the TCR. Uniform manifold approximation and projection (UMAP) analysis of single-cell RNA-seq of these cells revealed broad expression of *CSF2* but with separate cell clusters showing high expression of *IL17A* versus *IFNG*. In flow-chamber assays, we found that chemokine ligands for CCR5, CCR6, and CXCR3 could be found on the luminal surface of activated endothelial cells and each of these receptors could contribute to firm arrest but had no role in TEM. By contrast, the CCR2 ligands CCL2, CCL7 and CCL8 (also a ligand for CCR5), although secreted from both luminal and basal sides of endothelial cells, failed to bind to these cells’ luminal surfaces - and CCR2 did not contribute to arrest but had a dedicated role in TEM. Using a CCL2-CXCL9 chimeric chemokine, we found that surface-bound CCL2 could induce CCR2-dependent arrest, but at the cost of inhibiting TEM. Together, our findings establish the ability to extravasate efficiently as a capability of resting type 17 cells with pathogenic potential and demonstrate that this extravasation depends both on redundant activities of chemokine receptors at the step of firm arrest and a dedicated role for CCR2 in TEM that derives not from intrinsic differences in receptor signaling but from selective localization of chemokine ligands. During extravasation, arrest and TEM are mutually exclusive activities of chemokine receptors and their ligands - whereas arrest requires chemokine bound to EC surfaces, our data suggest that TEM requires a transendothelial gradient of chemokine created by removal of non-bound, luminal chemokine by vascular flow.

## Results

### CCR2 identifies pathogenic human type 17 cells

To understand the mechanisms used by bona fide human memory type 17 Th cells, for which we and others have found CCR6 can be used as a signature marker ^28, 29^, to traffic into inflamed tissue we took advantage of insights from our studies of CD8α^+^ MAIT cells, where we identified an important role for CCR2 ^27^. Consequently, we focused on resting, non-regulatory CD4^+^ cells that co-express CCR6 and CCR2. Because we found that CCR2^+^ cells within the CCR6^+^ population were generally CCR6^high^, for purposes of comparison we separated CD4^+^ cells from healthy donors into naïve and memory-phenotype populations and further separated HLA-DR^-^CD25^-/low^CD127^+^ memory-phenotype cells into those that were CCR6^-^CCR2^-^, CCR6^low^CCR2^-^, CCR6^high^CCR2^-^ and CCR6^+^CCR2^+^ (Fig. 1 A). It is important to note that for the data presented below we relied on flow cytometry and FACS using the panel of antibodies to identify and purify cell subsets as described in materials and methods. Particularly for CCR2, in which positive and negative cells do not form discreet populations, “negative” cells may nonetheless express low numbers of receptors.

**Figure 1.**
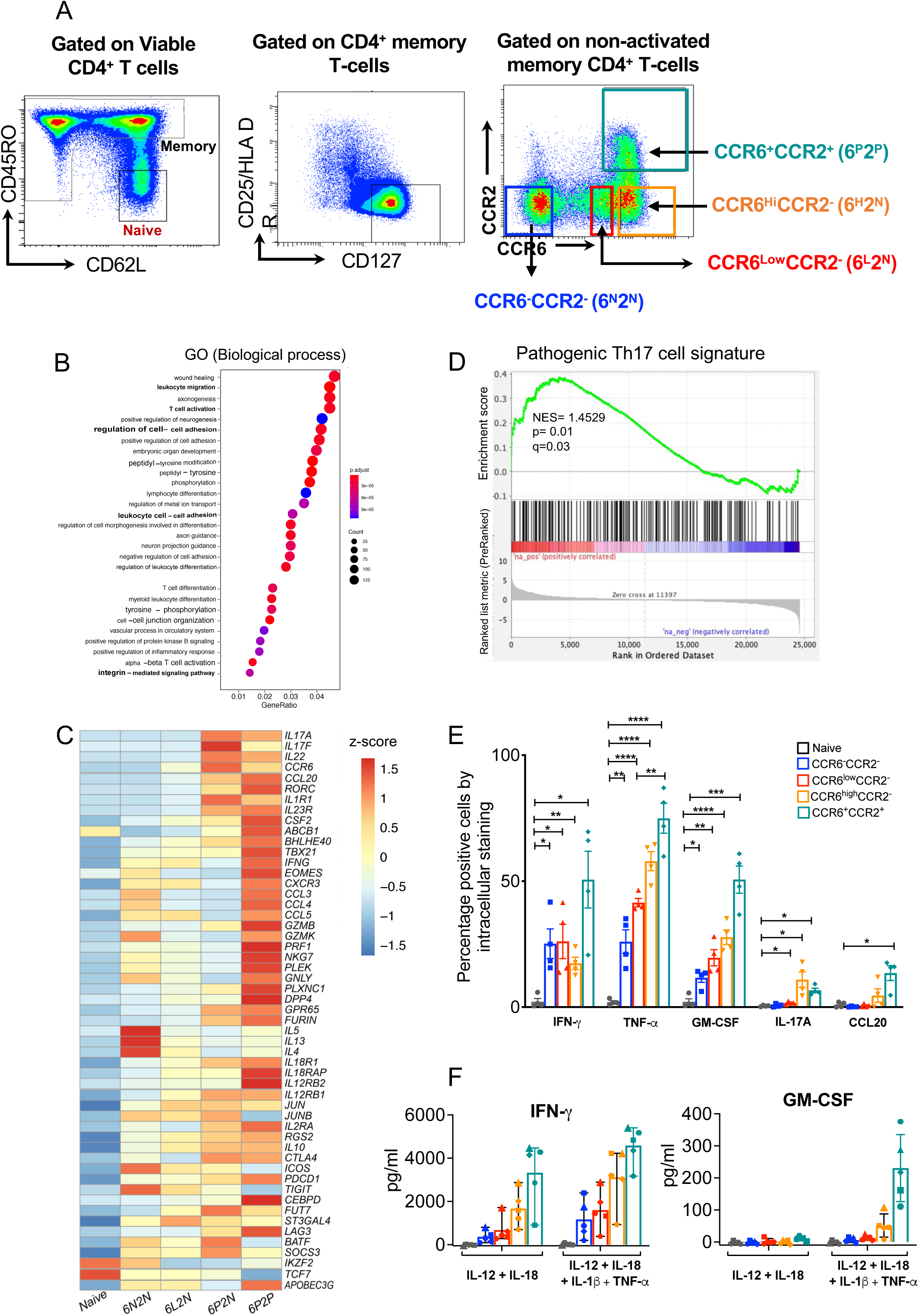
CCR2 identifies type 17 cells with a pathogenic profile. (A) FACS strategy to identify Naïve, CCR6^-^CCR2^-^ (6N2N), CCR6^low^CCR2^-^ (6L2N), CCR6^high^CCR2^-^ (6H2N), and CCR6^+^CCR2^+^ (6P2P), CD4 T cell subsets. (B) Gene Ontology (GO) biological process gene set analysis of top differentially expressed genes in CCR6^+^CCR2^+^ vs CCR6^-^CCR2^-^ cells. The size of the dots represent the number of genes in the significant DE gene list associated with the GO term and the color of the dots represent the adjusted P values. (C) Heatmap from bulk RNA-seq showing the expression levels of genes (rows) associated with the pathogenic Th17 phenotype in activated cells. Color coding reflects standardized gene expression values (z-scores). (D) GSEA for genes differentially expressed between activated CCR6^+^CCR2^+^ vs CCR6^-^ CCR2^-^ cells using a molecular signature gene set for pathogenic Th17 cells. (E) FACS-purified cells were activated ex-vivo and stained for indicated cytokines/chemokine as described in Materials and methods. In each case, staining with an isotype-matched control antibody was used to determine the percentage positive cells. Data are averaged from five donors and bars indicate means +/− SEM. p values were calculated using paired Student’s t tests, *, p<0.05; **, p<0.01; ***, p<0.001. (F) FACS-purified cells were stimulated ex-vivo with the indicated cytokines and culture supernatants collected 3 days post stimulation were assayed for GM-CSF and IFN𝛾 by ELISA. Data points are values from five donors and bars indicate means +/− SEM.

We performed RNA-seq on the FACS-purified cells from two donors. Gene Ontology (GO) analysis of the genes differentially expressed in CCR6^+^CCR2^+^ vs. CCR6^-^CCR2^-^ cells showed enrichment in genes associated with leukocyte migration, adhesion, and differentiation (Fig. 1 B). Type 17 genes such as *IL17A*, *IL17F, IL22, CCL20* and *RORC* were more highly expressed in the CCR6-positive subsets, particularly in the CCR6^high^CCR2^-^ and CCR6^+^CCR2^+^ cells (Fig. 1 C), which is consistent with what we and others have shown previously ^28, 29^. We noted that as compared with the other CCR6-expressing subsets, the CCR6^+^CCR2^+^ cells showed higher expression of genes associated with pathogenicity, based on mouse models of autoimmune disease and other data, such as *RORC*, *IL23R*, *TBX21, IFNG, BHLHE40*, *CSF2* (encoding GM-CSF)*, IL1R1* and *ABCB1* ^15, 16, 30^. Further, gene set enrichment analysis (GSEA) of the RNA-seq data from CCR6^+^CCR2^+^ vs. CCR6^-^CCR2^-^ cells showed a significant enrichment score against a gene set based on mouse data identifying a signature of pathogenic vs. non-pathogenic Th17 cells ^15^ (Fig. 1 D).

We next inspected expression of pro-inflammatory proteins of interest among chemokine receptors and cytokines. As shown in Fig. S1 A, the CCR6^+^CCR2^+^ cells showed, in addition to CCR6 and CCR2, increased expression of the inflammation-associated receptors CXCR6, CCR5 and CXCR3. Expression of the mRNAs for these receptors generally matched the pattern of surface staining (Fig. S1 B). Intracellular staining for cytokines in cells activated with PMA and ionomycin *ex vivo* (Fig.1 F) also matched the RNA-seq data. The CCR6^+^CCR2^+^ cells contained the highest percentages staining for the pathogenic cytokines IFN-γ, GM-CSF and TNF-α, but not for IL-17A. We also considered, given our earlier data suggesting a “first-responder” phenotype for CD4^+^CCR2^+^ cells ^22^ whether the CCR6^+^CCR2^+^ cells might produce effector cytokines after exposure to an inflammatory milieu in the absence of cognate antigen. As shown in Fig.1 E, among the subgroups of CD4^+^ T cells, the CCR6^+^CCR2^+^ cells secreted the most IFN-γ and GM-CSF after culturing with combinations of inflammatory cytokines, similar to the results following pharmacologic activation. Together, the data suggest that within the CCR6^+^, type 17 memory population, CCR2 marks cells with increased pathogenic potential.

### Single-cell analysis of CCR6^+^CCR2^+^ Th cells reveals heterogeneity in expression patterns for pathogenicity-associated genes

Heterogeneity of functional importance has been described for type 17 cells in mouse models of disease ^31, 32^ and we investigated heterogeneity among the CCR6^+^CCR2^+^ cells using the broad and unbiased approach of single-cell RNA-seq on freshly isolated CCR6^+^CCR2^+^ cells from two donors after activation with PMA and ionomycin. For purposes of comparison in the trajectory analysis (see below), we included non-selected CD4^+^ memory cells (Fig. 2 A). Data analysis for donors 1 and 2 are displayed in Figures 2 and S2, respectively. Uniform manifold approximation and projection (UMAP) plots were generated to create four clusters (Fig. 2 B and Fig. S2 A). Cluster heat maps (Fig. 2 C and Fig. S2 B) and dot plots (Fig. 2 D and Fig. S2 C) showed that *CSF2* is expressed broadly among the clusters, with a separation of cells with highest expression of the type 17 genes *IL17A*, *RORC*, *CCL20* and *IL22* from those with highest expression of the type 1 genes *IFNG*, *CCL3* and *CCL4* along with genes important for cytotoxicity, such as *PRF1, GZMK and GZMB*. Additional genes of interest include *IL1R*, which clusters with type 17 cytokines, *EOMES*, which clusters with type 1 cytokines/chemokines, and *IL23R* and *TBX21*, which show broad distributions. Feature scatter plots provide additional granularity, showing, for example in Fig. 2 E and Fig. S2 D, that the signal within cluster 1 for the type 1 gene *CXCR3* is lower in the region with highest expression of type 17 cytokines.

**Figure 2.**
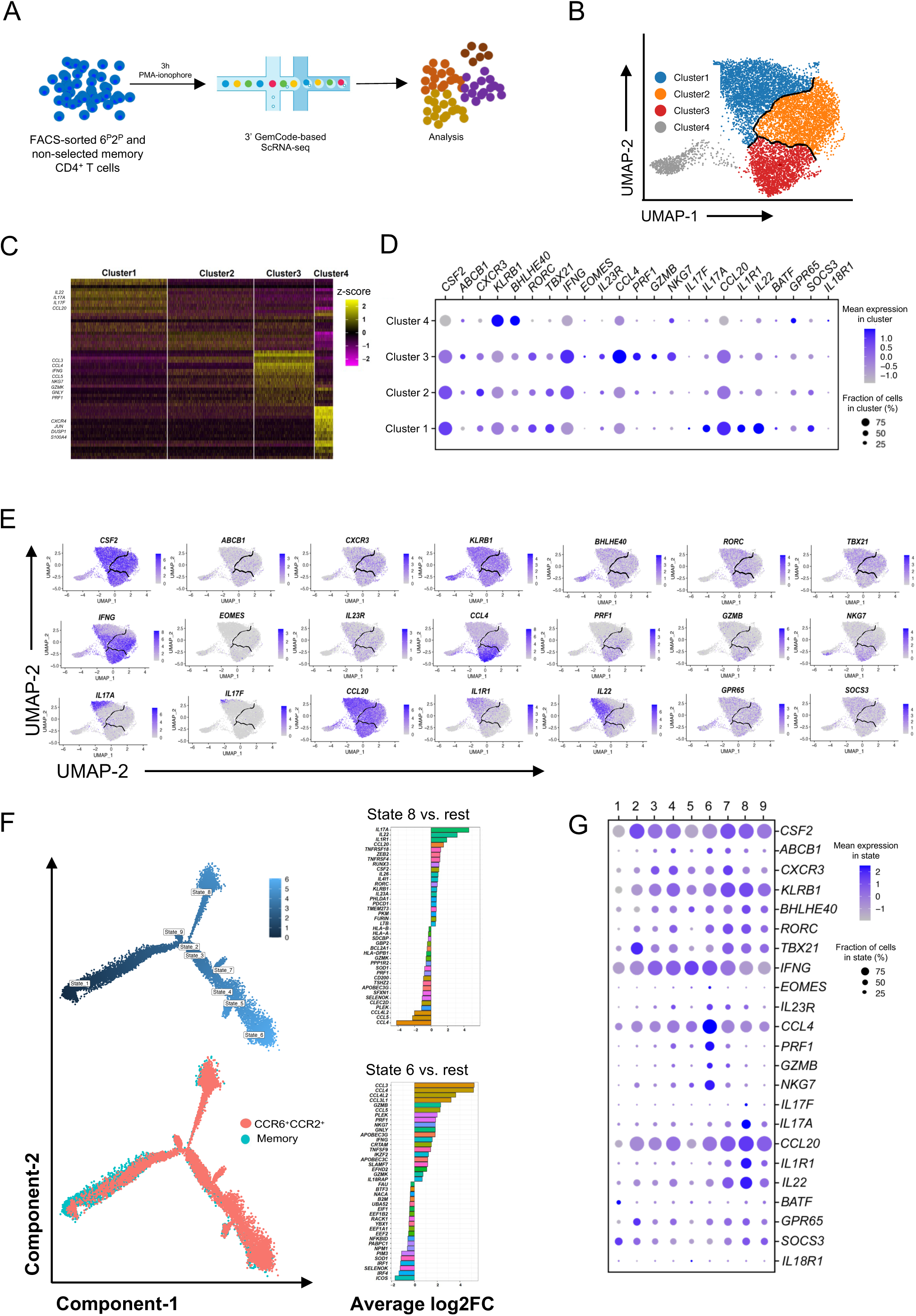
CCR6^+^CCR2^+^CD4+ T cells separate into clusters expressing *IL17A/F* vs. *IFNG*. (A) Schematic representation for single-cell analysis of FACS-purified cells. (B) UMAP projection and clustering from scRNA-seq profiles of 8,907 CCR6^+^CCR2^+^ cells from Donor 1 where each dot represents a single cell, with color codes of clusters defining cells with similar transcriptional profiles. (C) Heatmap showing the relative expression across all cells for the top upregulated and downregulated genes in each cluster compared to all other clusters. Rows represent genes and columns represent cells grouped by cluster, and some marker genes for clusters 1, 3 and 4 are listed on the left. Color coding reflects standardized gene expression values (z-scores). (D) Dot plot displaying gene expression for a subset of the genes displayed in (C). (E) Feature plots of expression levels of selected genes in cells in the UMAP scatter plot. Cluster borders are marked in black. (F) Pseudotime trajectories of single cell transcriptomic data showing states (upper left) and with cells colored according to their origin as either total unselected memory or CCR6^+^CCR2^+^ cells (bottom left). Right panel shows average log2 fold changes of genes with greatest differential expression in state 8 (upper right) or state 6 (bottom right) versus the remaining cells. (G) Dot plot shows the expression of selected genes in each state.

The trajectory analysis of CCR6^+^CCR2^+^ cells combined with non-selected memory cells formed three major branches with the non-selected memory cells mostly located in branches distinct from those dominated by the CCR6^+^CCR2^+^ cells (Fig. 2 F, left and Fig. S2 E). Consistent with the UMAP analysis, the gene bar plots for states 8 and 6 (Fig. 2 F and Fig. S2 E) and the state-wise expression dot plots (Fig. 2 G and Fig. S2 F) show separation of cells with highest expression of type 1 genes from cells with highest expression of type 17 genes, and with *CSF2* expression widely distributed. Together, these data suggest that the CCR6^+^CCR2^+^ subset contains highly differentiated memory cells that express a diverse collection of regulators and effectors of pathogenicity in distinct patterns, the highest expression of *RORC*, *IL1R1* and genes for type 17 cytokines in different cells than those expressing the highest levels *EOMES* and *IFNG*, and expression of *CSF2* and *TBX21* being readily detectable across both types of cells.

### CCR6^+^CCR2^+^CD4^+^ T cells are the cells migrating most efficiently across inflamed endothelium

We used flow chambers with cytokine-activated human umbilical vein endothelial cells (HUVECs) to study the trafficking behavior of the CCR6^+^CCR2^+^ cells as compared with the other CD4^+^ subsets. We set shear stress at 0.75 dyne/cm^2^ for four minutes to allow for T cell accumulation, after which the shear stress was increased to 3 dyne/cm^2^ for an additional 16 minutes. Data were collected by video microscopy for subsequent analysis and are displayed as cells per field that demonstrated the sequential steps of rolling, firm arrest, and transendothelial migration (TEM). Given the requirements for rolling before arrest and arrest before TEM, ratios of numbers of arrested cells/rolling cells and transmigrating cells/arrested cells are shown to evaluate the steps of arrest and TEM independently.

As compared with memory cells, fewer naïve cells were able to roll and virtually none were able to arrest and undergo TEM. Within the memory population there was a general increase in numbers of cells rolling and arresting and in the percentages of rolling cells arresting from the CCR6^-^CCR2^-^ to CCR6^low^CCR2^-^ to CCR6^high^CCR2^-^ to CCR6^+^CCR2^+^ cells, and only the CCR6^+^CCR2^+^ cells were able to undergo TEM (Fig. 3 A).

**Figure 3.**
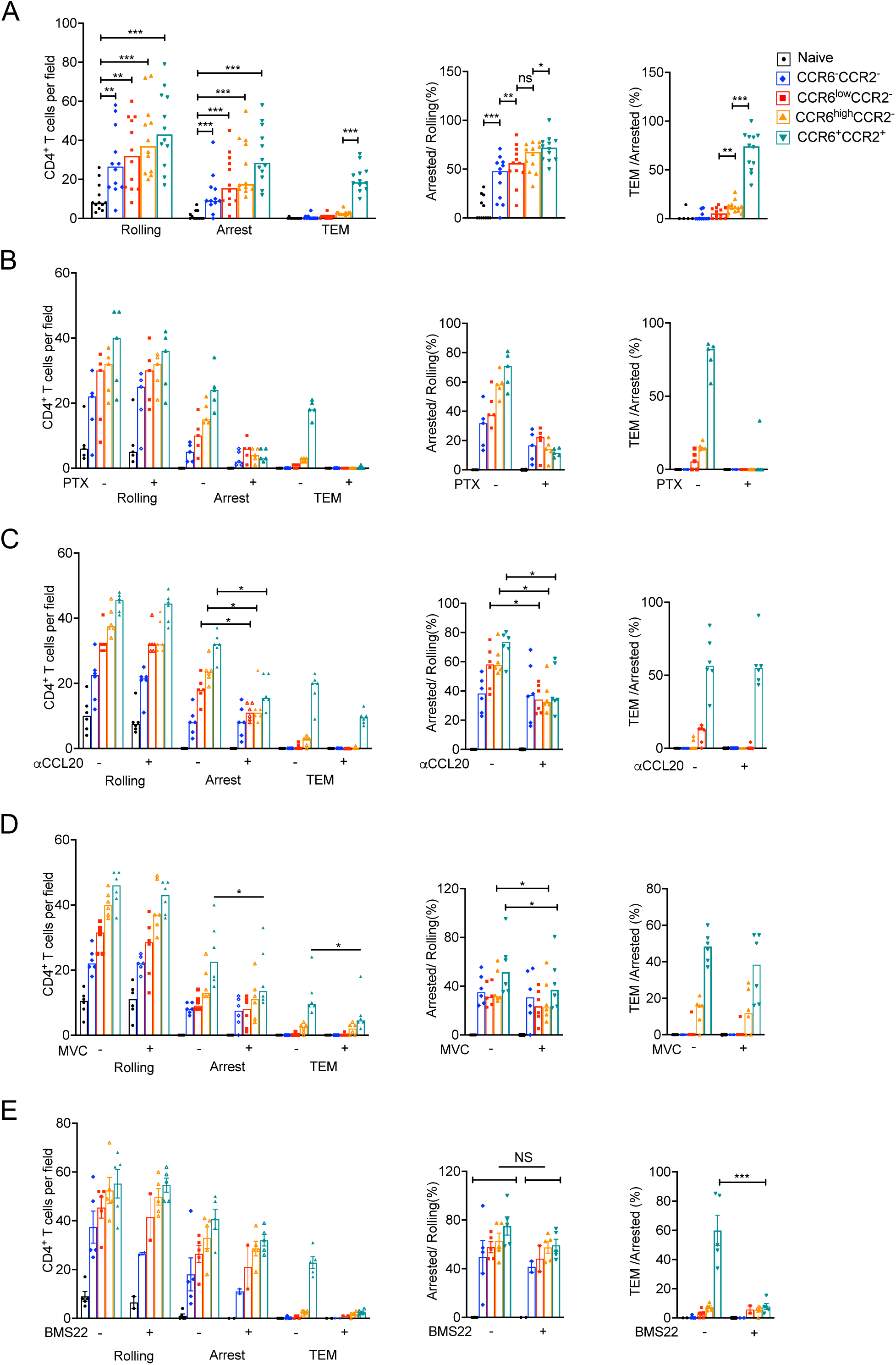
CCR6^+^CCR2^+^ are best at TEM. (A) Numbers of cells rolling, arrested and transmigrated on TNF-𝛼-activated HUVECs for CD4^+^ T cell subsets. Middle panel shows arrested cells as a percentage of cells rolling and in the right panel transmigrated cells as a percentage of cells arresting. Each symbol shows data from cells from one of 12 donors and bars indicate medians. (B) Data for T cells either untreated or treated with pertussis toxin. Data are from six donors and displayed as in (A). (C) Data for T cells in flow chambers using HUVECs pretreated with IgG control or anti-CCL20 antibody. T cells are from six donors and data are displayed as in A. (D and E) Data for T cells either untreated or treated with CCR5 antagonist, Maraviroc (D), or CCR2 antagonist, BMS22 (E). Data are displayed as in (A) and are from six (D) and five (E) donors, except that in two of the five experiments in (E) all the T cell subsets were treated with BMS22 whereas in the three additional experiments only the CCR6^high^CCR2^-^ and CCR6^+^CCR2^+^ cells were treated with BMS22. p values in (A), (C), (D) were calculated using Wilcoxon matched pairs signed rank test and in (E), using paired Student’s t tests. *, p<0.05; **, p<0.01; ***, p<0.001

Because chemokine receptor activation contributes variably to T cell arrest ^33^, we tested the receptors’ roles on our cells using pertussis toxin, which inhibits G_i/o_ proteins, the principal mediators of chemokine receptor signals. Pertussis toxin had no effect on rolling, diminished arrest by 50%-90% depending on the cell subset, and completely blocked TEM of the CCR6^+^CCR2^+^ cells (Fig. 3 B). To address the roles of individual chemokine receptors in these processes, we blocked specific receptors that we selected based both on the frequencies of their expression on the memory cell subsets and the levels of induction of chemokine-encoding mRNAs in the TNF-α-activated HUVECs (Fig. S3 A). Blocking CCR6 activity using a neutralizing antibody against CCL20 led to a significant reduction in arrest of CCR6-expressing cells, but no effect on the percentage of arresting CCR6^+^CCR2^+^ cells undergoing TEM (Fig. 3 C). Blocking CCR5 with the antagonist maraviroc resulted in a modest reduction in arrest, which reached statistical significance in the CCR6^high^CCR2^-^ and CCR6^+^CCR2^+^ cells (Fig. 3 D). These cells had shown the greatest sensitivity to pertussis toxin-mediated inhibition of arrest and had the highest frequencies of CCR5 expression (Fig. 1 B and S1 B). Just as after neutralizing CCL20, blocking CCR5 had no effect on TEM of the CCR6^+^CCR2^+^ cells (Fig. 3 D). By contrast, inhibiting CCR2 using the antagonist BMS22 had no effect on arrest but almost eliminated TEM by the CCR6^+^CCR2^+^ cells (Fig. 3 E), demonstrating a dedicated role for CCR2 in the final step of extravasation.

As in our previous studies on MAIT cells ^27^, these experiments were limited to endothelial cells activated by TNF-α. To broaden the conditions in the flow chamber assay and expand the collection of chemokines and receptors available to contribute to extravasation, we activated HUVECs with both TNF-α and IFN-γ. Combining these two cytokines increased expression of CCL5, the CCR5 ligand, induced expression of CCL7 and CCL8, additional ligands for CCR2, and significantly boosted expression of the three CXCR3 ligands, CXCL9, CXCL10 and CXCL11 (Fig. S3 B). The addition of IFN-γ led to increases in arrest by the memory cells with the largest and smallest increases in the CCR6^-^CCR2^-^ and CCR6^+^CCR2^+^ cells, respectively. There was no effect on rolling or on TEM *per se* (Fig. 4 A). Having already established roles for CCR6 and CCR5 in arrest and given the high-level induction of CXCR3 ligands with the addition of IFN-γ, we tested the effect of a CXCR3 antagonist in the flow chamber assay. As for CCR6 and CCR5, blocking CXCR3 diminished arrest, particularly on the CCR6^-^CCR2^-^ and CCR6^low^CCR2^-^ cells, which show a combination of no or little CCR6 and high expression of CXCR3 (Fig. 4 B). Taken together, our data show redundant but context-dependent activities of CCR6, CCR5 and CXCR3 in memory T cells limited to arrest, and a non-redundant role for CCR2 in TEM.

**Figure 4.**
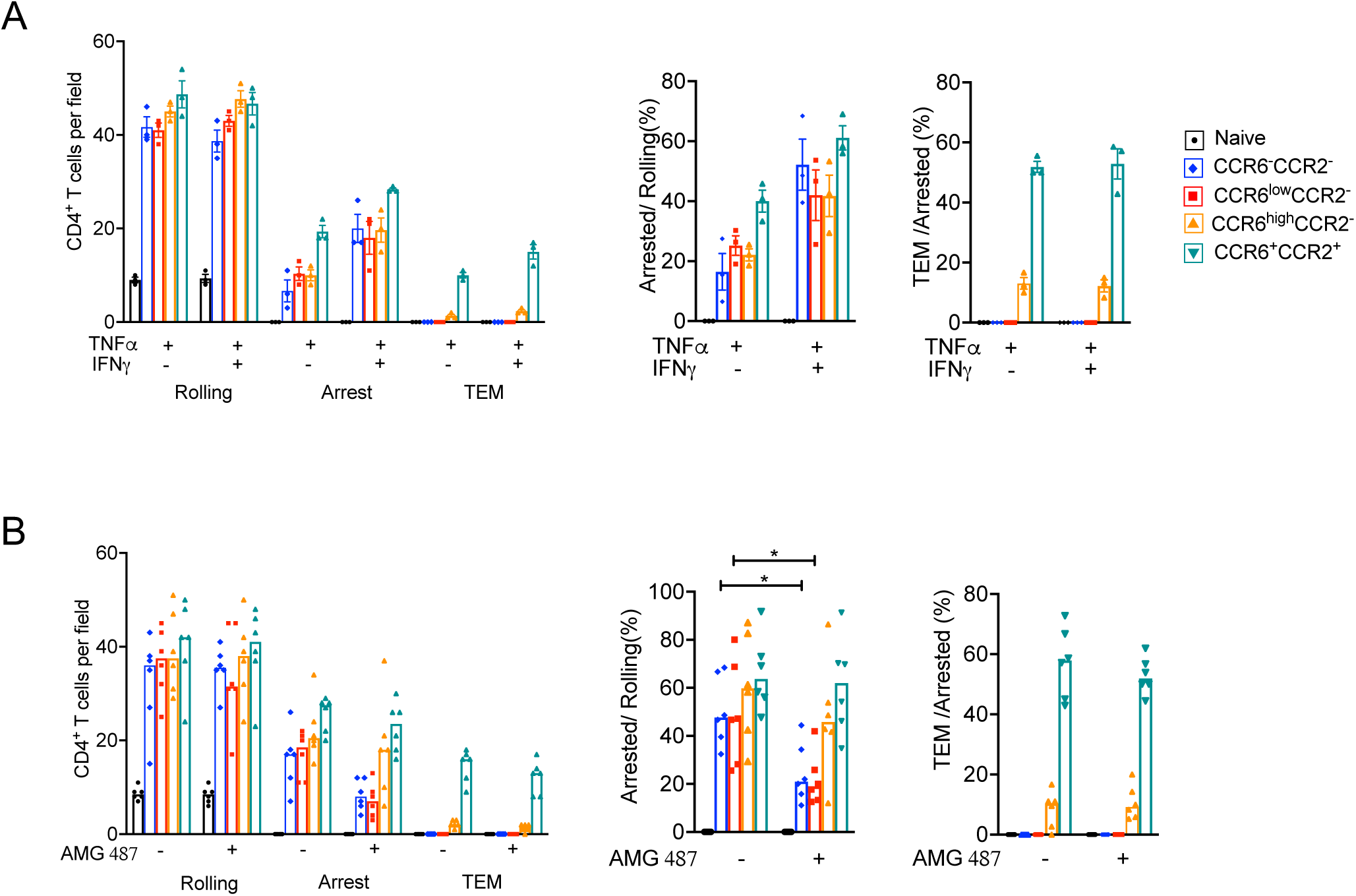
More cells arrest on TNF-𝛼 + IFN-𝛾-activated HUVECs. (A) Numbers of cells from CD4^+^ T cell subsets rolling, arrested and transmigrated on HUVECs activated with TNF-𝛼 alone or TNF-𝛼 + IFN-𝛾. Middle panel shows arrested cells as a percentage of cells rolling and in the right panel transmigrated cells as a percentage of cells arresting. Each symbol shows data from cells from one of three donors and bars indicate means +/− SEM. (B) Data for T cells either untreated or treated with CXCR3 antagonist, AMG487. Data are from six donors and displayed as in (A), except bars indicate medians. p values were calculated using Wilcoxon matched pairs signed rank test. *, p<0.05; **, p<0.01; ***, p<0.001

### Chemokine localization restricts and preserves CCR2 for a role in TEM

To understand the basis for the activities of the chemokine receptors in the steps of extravasation we investigated the localization of chemokines produced by activated HUVECs. We first measured chemokine found in the culture medium (supernatant) versus cell-associated for CCL20 and CCL2 after treating HUVECs with TNF-α and/or IFN-γ. The distributions of these chemokines were markedly different independent of the cytokine inducers. CCL20 was found more cell-associated than in the supernatant, whereas, comparatively, CCL2 was found more in the supernatant than associated with cells (Fig. S3 C). The patterns for CCL7 and CCL8 were like that for CCL2 after activating HUVECs with TNF-α plus IFN-γ (Fig. S3 C). We next tested whether secretion of these chemokines by the HUVECs was polarized by culturing HUVECs that were untreated or activated with TNF-α plus IFN-γ on transwell membranes and measuring chemokines in the upper and lower wells. CCL2, CCL7 and CCL8 were secreted into both upper and lower wells (Fig. 5 A).

**Figure 5.**
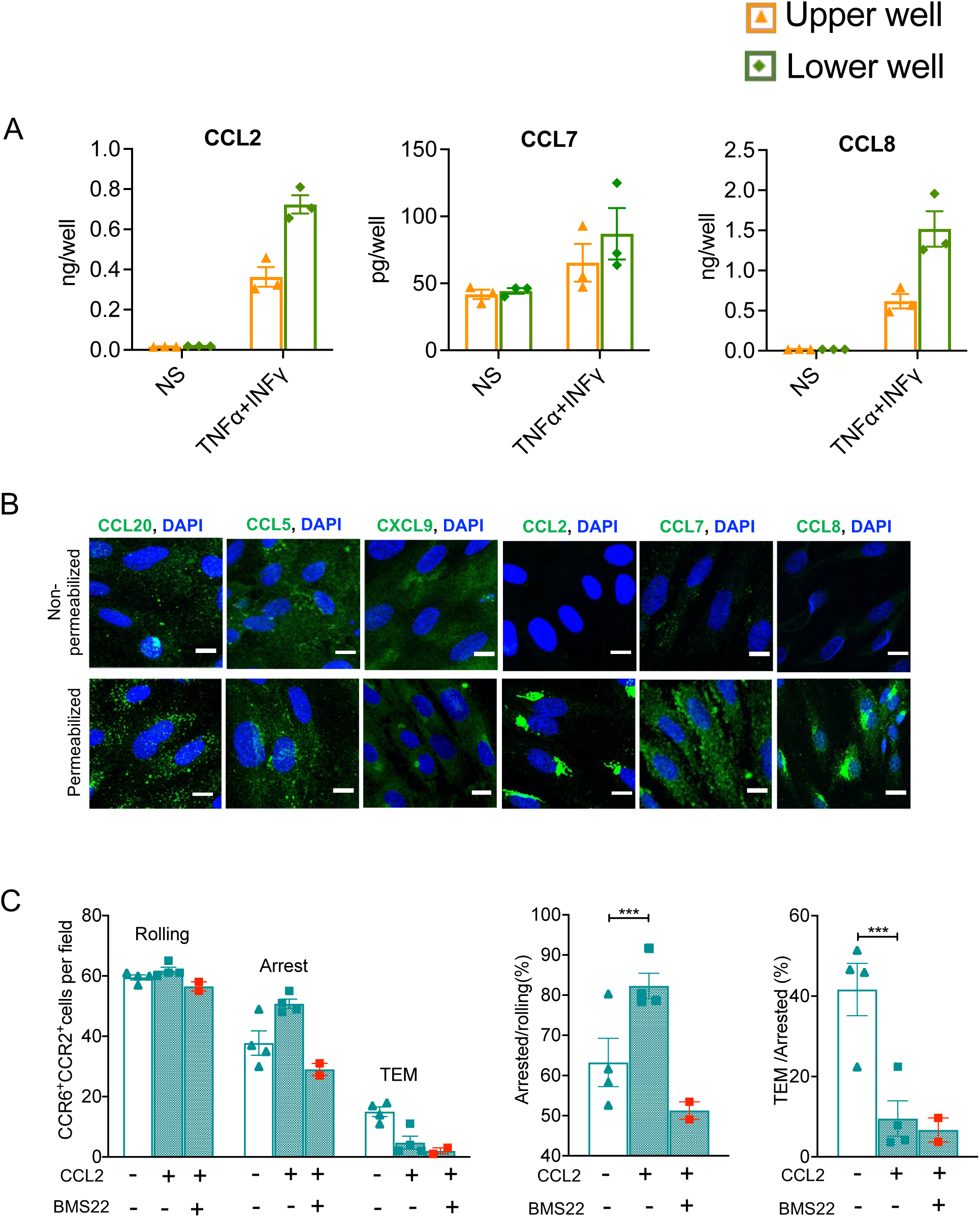
CCR2 ligands are secreted but do not bind to HUVECs. (A) Culture supernatants collected from the upper and lower wells of transwells with HUVEC monolayers were assayed for CCL2, CCL7 and CCL8 by ELISA. Data are from three experiments and bars indicate means +/− SEM. (B) Confocal microscopy images at ×40 magnification of non-permeabilized and permeabilized TNF-*α* + IFN-*γ*-stimulated HUVECs immunostained for CCL20, CCL5, CXCL9, CCL2, CCL7 or CCL8 (green) along with nuclear staining (blue) as described in Material and methods. The scale bars indicate 10 µm. Images are representative of three experiments. (C) CCR6^+^CCR2^+^ cells were either left untreated or treated with CCL2 (100 ng/ml) just before and for 4 min after adding to the flow chambers, and CCL2-treated cells were used with or without treatment with the CCR2 antagonist, BMS22. Data points in the left panel show numbers of cells rolling, arrested and transmigrated, while the middle panel shows arrested cells as a percentage of cells rolling and the right panel shows transmigrated cells as a percentage of cells arresting. Each symbol shows data from cells from one donor, with four experiments using CCL2 and two experiments including the additional treatment with BMS22. Bars indicate means +/− SEM. p values were calculated using paired Student’s t tests, *, p<0.05; **, p<0.01; ***, p<0.001.

To localize chemokines on and/or in endothelial cells, we used fluorophore-conjugated antibodies to stain HUVECs treated with TNF-α and IFN-γ before and after permeabilizing the cells. Their different staining patterns notwithstanding, ligands for CCR6 (CCL20), CCR5 (CCL5) and CXCR3 (CXCL9) could all be identified on both the surface and within the HUVECs by confocal microscopy, whereas the ligand for CCR2 (CCL2) could only be detected within the cells (Fig. 5 B). We also evaluated the other CCR2 ligands, CCL7 and CCL8, and found that they, too, are secreted but could not be detected on the surface of HUVECs (Fig. 5 B and S3 C). Additionally, we tested the binding of CCL2 and CCL5 to untreated and TNF-α-activated HUVECs using synthesized proteins. These chemokines were biotinylated particularly to detect their binding (using streptavidin-phycoerythrin) to the surface of activated HUVECs, where anti-chemokine antibodies would be expected to detect also endogenous chemokines. As for our findings for the endogenous chemokines, synthesized CCL5, but not CCL2, bound to the surface of HUVECs. Results were the same whether the HUVECs were untreated or activated with TNF-α (Fig. S3 D). Taken together, these results provided a straightforward explanation for the restricted activity of CCR2 and for the activities of CCR6, CCR5 and CXCR3 in arrest.

### Triggering arrest and TEM are mutually exclusive activities

The dedicated role for CCR2 in TEM in the absence of surface-bound ligands led us to ask if CCR2 were uniquely constituted for TEM or if the receptor could mediate arrest if activated in the appropriate context by extracellular ligand. We tested CCR2’s capabilities by adding CCL2 to CCR6^+^CCR2^+^ cells immediately before loading them into the flow chamber. Under these conditions, CCL2 did, in fact, increase the cells’ arrest, and this increase was reversed by blocking CCR2. It was of interest that while adding CCL2 enhanced arrest, TEM was suppressed (Fig. 5 C). This latter effect might be expected if the concentration of CCL2 were sufficient to desensitize/internalize CCR2. However, preincubating cells with CCL2 at the concentration used in the flow chamber assay (100 ng/ml) did not lead to receptor internalization, which required concentrations that were a significantly higher (Fig. S4).

While these experiments demonstrated that CCR2 is capable of triggering arrest in the presence of high concentrations of soluble ligand, we sought to produce a more physiologically appropriate CCR2 “arresting” ligand by having endothelial cells secrete a form of CCL2 that would bind to surface glycosaminoglycans, analogous to ligands for other chemokine receptors. This was done by transducing HUVECs with a lentiviral vector containing sequences encoding a chimeric chemokine consisting of full-length human CCL2 followed by carboxy-terminal residues 74-103 of human CXCL9 (Fig. 6A, top). These carboxy-terminal sequences of CXCL9 are highly basic ^34, 35^ and have been shown to bind glycosaminoglycans ^34, 35^, and we demonstrated that HUVECs transduced with the chimeric sequences now showed surface staining with the antibody directed against CCL2 (Fig. 6A, bottom). We purified control-transduced and chimera-transduced GFP^+^ HUVECs by FACS, and after activating them with TNF-α used them in flow chamber assays with CCR6^+^CCR2^+^ cells. We found increased numbers of CCR6^+^CCR2^+^ cells arresting on the chimera-expressing HUVECs, and these increases were reversed by inhibiting CCR2. Additionally, as in our findings using an immediate pre-incubation with CCL2-containing medium, the surface-bound CCL2-CXCL9 chimera effectively blocked TEM (Fig. 6 B). Taken together, these data suggest ligand localization rather than intrinsic properties determine a chemokine receptor’s role in extravasation, and that on these cells roles for receptors and their ligands in arrest and TEM are mutually exclusive.

**Figure 6.**
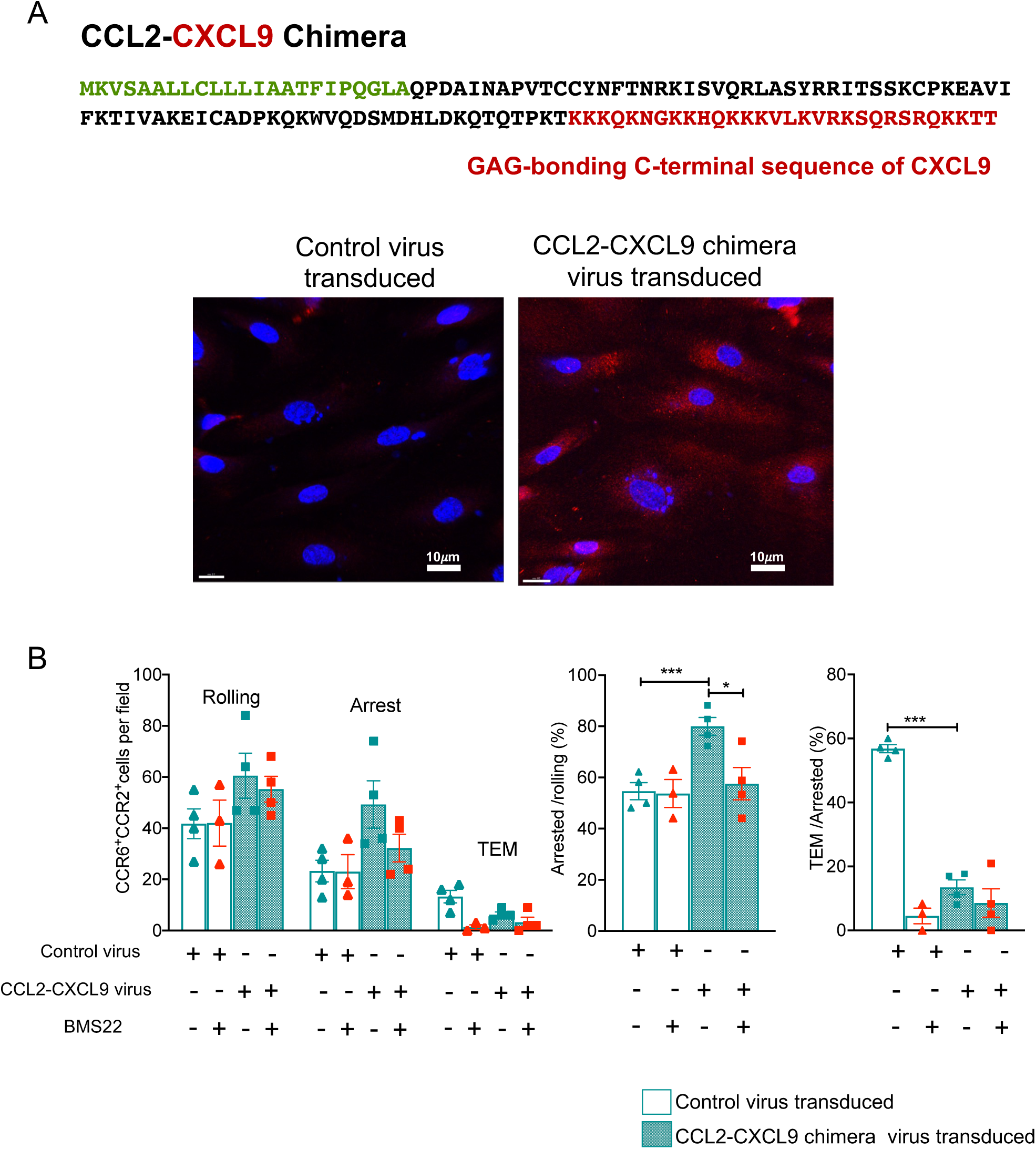
Surface-bound CCL2 can induce CCR2-mediated arrest. (A) CCL2- CXCL9 chimera sequence with signal peptide sequence shown in green, CCL2- sequence in black and C-terminal GAG-binding sequence of CXCL9 in red (top). Confocal microscopy images of control or CCL2-CXCL9 chimera transduced, TNF-*α*-stimulated HUVECs immunostained for CCL2 (red) and DAPI (blue) as described in Materials and methods (bottom). (B) Numbers of CCR6^+^CCR2^+^ cells, either untreated or treated with the CCR2 antagonist, BMS22, rolling, arrested and transmigrated on TNF-𝛼-activated HUVECs transduced with either control virus or with virus encoding the CCL2-CXCL9 chimera. Middle panel shows arrested cells as a percentage of cells rolling and in the right panel transmigrated cells as a percentage of cells arresting. Each symbol shows data for cells from one donor, with cells from four donors in each treatment group except for the BMS22-treated cells with control virus-transduced HUVECs, where cells from three donors were used. Bars indicate means +/− SEM. p values were calculated using paired t tests, *, p<0.05; **, p<0.01; ***, p<0.001.

## Discussion

In our previous work we found that CCR6 is expressed on all human T cells capable of making IL-17A ^29^ and, in fact, CCR6 identifies a broad family of related “type 17” human Th cells that are particularly suited to barrier defense ^17^. We also described that CCR2 is co-expressed with other inflammation-associated chemokine receptors, including CXCR3, CCR4, CCR5 and CCR6 on highly differentiated, effector-capable human Th memory cells equipped to serve as early recruits to an immune response ^22^. In investigating other CCR2^+^ human T cells, we found that the receptor is detected on most MAIT cells, where it is co-expressed with a similar complement of chemokine receptors and has a non-redundant role in TEM ^27^. We reasoned that CCR2 might mark a pro-inflammatory subset of CCR6^+^/type 17 cells that have, as one of their functionalities, the ability to migrate efficiently into inflamed tissue. We found that the profile of gene and protein expression of CCR6^+^CCR2^+^ Th cells resembles what has been reported for pathogenic Th17 cells, and, consistent with co-expression of CXCR3, for cells for cells described, according to the experimental context, in mice or humans as “ex-Th17” or Th1-like or Th1* cells ^16, 28, 36, 37^.

In mouse models these cells mediate pathology in experimental autoimmune encephalomyelitis (EAE) ^31^, autoimmune colitis ^36^ and arthritis ^38^. In humans these cells have been strongly implicated in host defense against mycobacteria ^39–42^ and in the autoimmune diseases of multiple sclerosis ^43–45^ and rheumatoid arthritis ^46^. To manifest pathogenicity in mice these cells have been reported to require IL-23 ^15, 47, 48^ and the transcriptional regulators RORγt and T-bet ^36^, JunB and BATF ^49^, BHLHE40 ^50^, and EOMES ^51^. The critical effector cytokines that have been shown to mediate the pathogenic activity of these cells have included GM-CSF ^38, 48, 51–58^, IFN-γ ^36, 52^ and IL-17 ^55^. These cells express multiple granzymes that may be important for tissue damage ^59^, and also express a cytotoxic program ^39^. Surface proteins described on these cells in humans include CCR6 and CXCR3 ^28, 37, 46^ and MDR1/ABCB1 ^30, 46^.

We found that many of the relevant genes, including *IL23R*, *IL18R1*, *IL18RAP*, *RORC*, *TBX21*, *BHLHE40*, *EOMES*, *CSF2*, *IFNG*, *CXCR3*, *PRF1*, *GZMB*, *GZMK*, *GNLY*, *NKG7*, and *ABCB1* were enriched in the CCR6^+^CCR2^+^ vs. the CCR6^high^CCR2^-^ subgroups of human Th cells and the percentages of cells capable of producing GM-CSF and IFN-γ after pharmacologic activation *ex vivo* were similarly increased in the CCR6^+^CCR2^+^ vs. the CCR6^high^CCR2^-^ subgroups. The importance of CCR6^+^CCR2^+^ cells in host defense, particularly early during pathogen challenge, and as contributors to immunopathology would be enhanced if they were able to produce effector cytokines in response to an inflammatory environment in the absence of cognate antigens. In fact, we found that these cells were the CD4^+^ T cells that secreted the most IFN-γ and GM-CSF solely in response to combinations of inflammation-associated cytokines. Recent data in mice have suggested that this sort of cytokine-induced, bystander activation of CD4^+^ T cells can contribute to the immunopathology of EAE ^60^. CCR2 itself has been shown to be required in mice for EAE-mediated disease due to the receptor’s critical role in monocyte migration ^61, 62^. CCR2 has also been reported to contribute to EAE by mediating recruitment to the central nervous system of a GM-CSF-producing subset of cells characterized as CCR6^-^CCR2^+^ ^56^, and CCR2^+^ Th cells were found to be enriched in the CSF of patients with multiple sclerosis during relapse ^63^. These latter cells had features, including the ability to produce IFN-γ in response to myelin basic protein, suggesting a role in pathogenesis ^63^.

Clustering of CCR6^+^CCR2^+^ cells from our single-cell RNA-seq analysis revealed differences in the distributions of mRNAs encoding proteins associated with pathogenicity. Of particular interest, within the major clusters, there was widespread expression of *CSF2*, whereas expression of *IL17A/F* and *IFNG* did not overlap. The distribution of expression of *IL1R1* matched that of *IL17A/F*, while expression of *CCL4*, *PRF1* and *EOMES* matched that of *IFNG*. The coincidence of expression of *IL1R1* and *IL17A/F* is consistent with the critical role for IL-1β in the differentiation of human Th17 cells ^28, 64^, in marking Th17 cells and enhancing their production of IL-17A ^65^ and, in combination with IL-12, enabling human CCR6^+^ Th cells to acquire a pathological phenotype ^66^. Co-localization of expression in the UMAP-generated display for *IFNG* with the chemokines *CCL3*, *4*, and *5*, which encode ligands for CCR5 and CCR1, is consistent with the established co-expression of these chemokines with IFN-γ ^67–69^. The expression of *PRF1*, encoding perforin, and/or genes encoding granzymes have been reported in Th cells, including Th17 cells ^70, 71^, that are implicated in autoimmunity ^72, 73^ and host defense ^71^, and that co-express EOMES ^72, 74^ and IFN-γ ^73^. Taken together, our data suggest that the CCR6^high^CCR2^-^ cells most resemble conventional type 17 cells, whereas the CCR6^+^CCR2^+^ Th subgroup is enriched in type 17 cells with pathogenic capabilities. Further, although expression of *CSF2* is a general property of the CCR6^+^CCR2^+^ Th cells, these cells tend to fall along a gradient between two polar phenotypes: 1) with features of IL-17A/F producing cells, which express IL-1R1, and 2) with type 1 features, including the production of IFN-γ and expression of *EOMES* and molecules important for cytotoxicity. The trajectory analysis suggested that these two subsets of CCR6^+^CCR2^+^ Th cells with pathogenic features separate over pseudotime from common precursors to form two branches of more differentiated cells.

The ability to migrate into inflamed tissue is integral to the function of effector-capable T cells. The resting memory-phenotype cells that we studied here would be available as early responders from the blood to reinforce resident memory cells in the event of a pathogen challenge or to contribute to immunopathology ^75^. Although there have been a few highly informative studies of roles for individual chemokine receptors on human T cells, these experiments used heterogeneous populations of T cells that were activated and expanded in vitro and presumably corresponded most closely to recently activated effector cells ^76, 77^. Work from our laboratory, including in the current studies, includes, as far as we are aware, the only investigations describing roles in extravasation for individual chemokine receptors on non-manipulated populations of highly differentiated memory T cells, which typically express multiple receptors ^27^. Within the human Th cell population, we show here activities of CCR5, CCR6, and CXCR3 in firm arrest. The activities for CCR5 and CCR6 only required activation of HUVECs with TNF-α, whereas the activity of CXCR3 depended on including IFN-γ to induce CXCR3 ligands, and for each of these receptors we demonstrated the display of a cognate chemokine on the HUVEC surfaces. None of these receptors could mediate TEM. The recruitment of cells that express CXCR3, the signature chemokine receptor for Th1 cells, in response to treatment of HUVECs with IFN-γ would support a positive feedback loop to amplify a type 1 response. The contributions of individual receptors to arrest among the Th cell subgroups were consistent with the subgroups’ patterns of receptor expression. For example, CXCR3 contributes the most to arrest of the CCR6^-^CCR2^-^ and CCR6^low^CCR2^-^ cells, which both express CXCR3 and have low levels or no expression of the other arrest-mediating receptors, CCR5 and CCR6. By contrast with these receptors, CCR2 was unable to contribute to firm arrest, but was the only receptor mediating significant TEM. This latter activity profile was associated with the secretion of the CCR2 ligands CCL2, CCL7 and CCL8 by HUVECs treated with TNF-α plus IFN-γ into the culture medium from both luminal and basal sides, but a failure of these chemokines to be retained on the HUVEC surfaces.

The display of chemokines on the surfaces of endothelial cells depends on binding to glycosaminoglycans (GAGs), primarily heparan sulfate ^78, 79^, and eliminating heparan sulfate diminishes leukocyte trafficking to lymph nodes ^80, 81^ and inflamed tissue ^82^. Although the complexity and heterogeneity of glycosaminoglycans has hindered the identification of the physiological moieties binding to chemokines, existing data suggest significant specificity in GAG-chemokine interactions, including for CCL5, CCL2, and CCL20 ^79, 83, 84^. Although CCL5 has a particularly high affinity for GAGs ^83^, CCL2 and CCL7 also bind ^79, 85–87^, and the published data do not explain why we could not detect CCL2 or the other CCR2 chemokines binding to HUVECs. Nonetheless, other investigators have also reported the absence of CCL2 binding to endothelial cells ^33, 88, 89^. Some possible reasons for these findings might include unappreciated post-translational modifications of the secreted CCL2, or CCL2 being secreted already complexed to GAGs, such as has been described for CCL3, 4, and 5 from CD8^+^ T cells ^90^, or competition between CCL2 and other chemokines or non-chemokine proteins for GAG binding. Our finding that not only endogenous, but also recombinant CCL2 failed to bind to both non-activated and activated HUVECs would not support the first two mechanisms, nor competition between CCL2 and other cytokine-induced chemokines as parsimonious explanations. Competition between the CCR2 ligands and non-chemokine GAG-binding proteins remains plausible. These data do not preclude the possibility that CCL2 or other CCR2 ligands may bind to the surfaces of some non- HUVEC endothelial cells *in vivo*, such as has been reported for HEV in mouse lymph nodes draining inflamed skin ^91^.

CCR2-mediated arrest after adding CCL2 to T cells immediately before they entered the flow chamber or, more interestingly, by using HUVECs displaying GAG-bound CCL2 as part of a CCL2-CXCL9 chimera, demonstrated that the failure of CCR2 to mediate arrest was solely due to the lack of sufficient ligand at the site and time of the initial T cell-HUVEC interactions. We showed that CCL2 and other CCR2 ligands are secreted from both sides of HUVECs. We presume that the dilution due to continuous flow on the luminal side coupled with basilar secretion establishes a transendothelial concentration difference that drives CCR2-dependent TEM. The inhibition of TEM from exposing cells to high concentrations of surface bound CCL2 on the luminal side would follow from the loss of this transendothelial difference.

It has been reported that the delivery of CCL2 to invasive filopodia at sites of T cell-endothelial cell contact at the luminal surface by intracellular vesicles is a mechanism for CCR2- dependent TEM of activated T cells ^33^. However, other studies support the importance of a transendothelial difference in ligand concentration for CCR2-mediated TEM of leukocytes ^76, 88, 89^. Additionally, there are data suggesting the preferential secretion of chemokines from the basal surface of endothelial cells ^76^, which, along with a gradient of heparan sulfate from the lumen to the basolateral surface of post-capillary venules, as has been described ^92^, would be expected to enhance a transendothelial difference. A study of ischemia/reperfusion-induced kidney injury in mice and humans reported the accumulation of CCL2 on microvascular basement membranes, which could also serve as a depot of subendothelial chemokine *in vivo* ^93^. Fig. 7 shows a cartoon illustrating our view of the steps and mediators of extravasation of the CCR6^+^CCR2^+^ cells.

**Figure 7.**
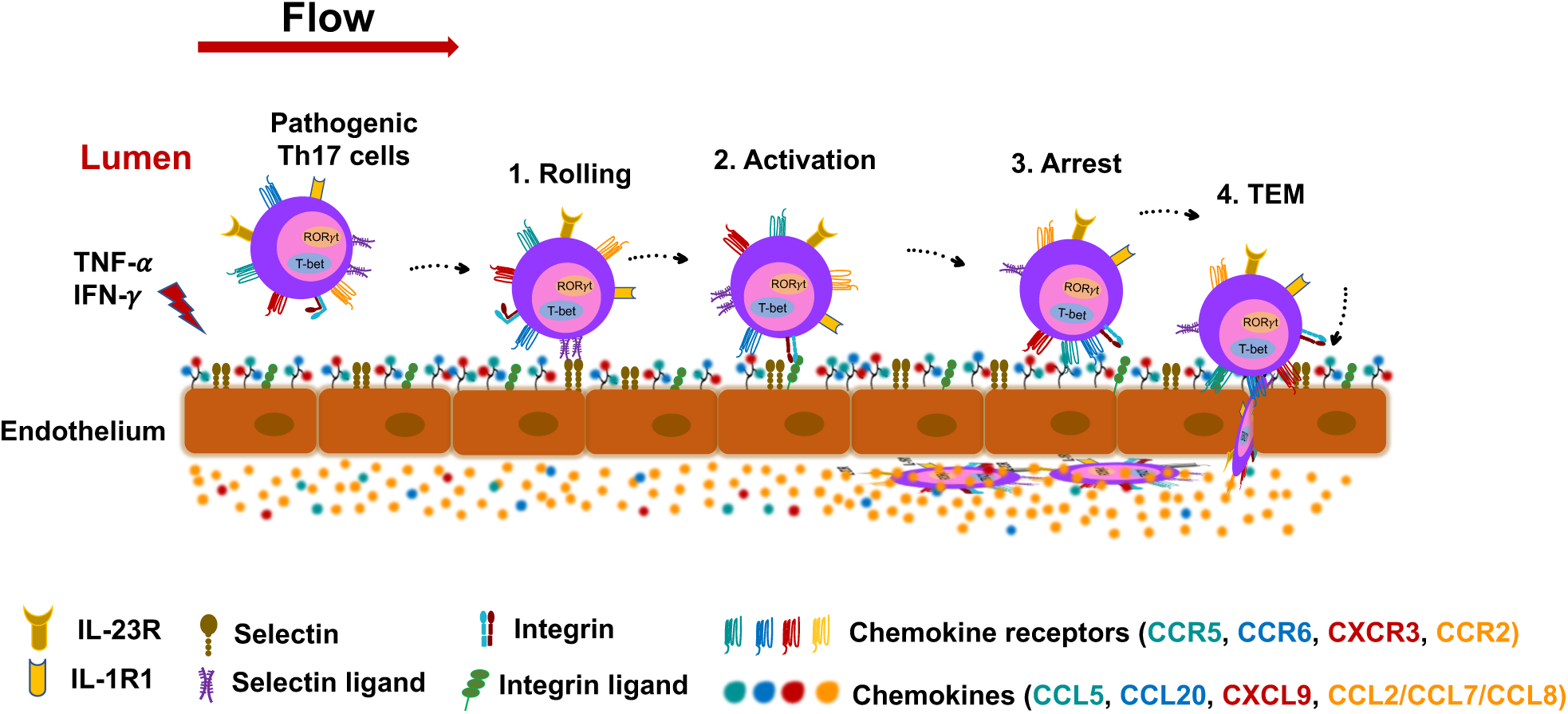
Extravasation of pathogenic human type 17 Th cells across inflamed endothelium. Inflamed endothelial cells upregulate expression of selectins and integrin ligands and secrete various chemokines. Selectin-selectin ligand interactions help capture cells and allow them to roll along the endothelium. For pathogenic Th17 cells, endothelial cell-bound chemokines can then stimulate CCR6, CCR5 and CXCR3 to activate integrins for binding to intercellular adhesion molecules and mediate firm arrest. Secreted chemokines that fail to adhere to endothelial cells, such as CCL2 and other CCR2 ligands, are diluted by blood flow to create a transendothelial gradient that induces TEM.

Our data on the mutually exclusive activities of chemokine receptors and their ligands in arrest versus TEM raise some additional issues. Firstly, although it is clear from experiments using Boyden Chamber assays under static conditions that leukocytes can migrate in response to a transendothelial difference in CCL2 concentration ^88, 95^, precisely how a leukocyte senses a chemokine gradient across the endothelial cell barrier remains uncertain. Consistent with our findings, elegant studies of the extravasation of mouse neutrophils *in vivo* have shown mutually exclusive roles for chemoattractants and/or chemoattractant receptors in arrest versus TEM ^94, 96^. However, one of these studies on neutrophils differed from our work in showing that although arrest and TEM were mediated by different chemokines, both ligands were acting through one receptor, CXCR2 ^94^ – a finding that could have been due to ligand-biased signaling (see below.) In these reports, the CXCR2 ligand(s) mediating TEM were produced by non-endothelial cells and transported and/or presented on endothelial cells by the atypical chemokine receptor, ACKR1, which was shown to localize preferentially at endothelial cell junctions ^94^. In our experiments, the activated endothelial cells themselves were the source of the CCR2 ligands mediating TEM, thereby obviating a requirement for ACKR1 in endothelial cell transport and/or presentation, and we found no evidence of CCR2 ligands localized at endothelial cell junctions. Nonetheless, it is possible that chemokine secreted on the abluminal side could be presented at the junctions by endothelial cell ACKR1 and form a gradient sensed by the leukocytes’ invading podosomes ^97^. In this regard, both CCL2 and CCL7 bind well to ACKR1 ^98^. Alternatively, leukocytes could similarly sense soluble gradients extending from the abluminal side into the endothelial cell junctions, which are opened at the luminal side by leukocyte-endothelial cell interactions prior to the initiation of TEM ^99^.

Secondly, the inhibition of TEM by the CCL2-CXCL9 chimera might suggest that T cell migration should likewise be inhibited by endothelial-cell bound chemokines such as CCL5, CCL20 and CXCL9, since moving away from the luminal surface could involve migrating in opposition to the concentration difference in chemokine. However, based on our and published data ^76^, like CCL2, other chemokines, including “arrest chemokines” that bind to endothelial cells, are also secreted at both luminal and basal sides of endothelial cells, minimizing the size of any transendothelial gradient that would oppose TEM. *In vitro* experiments on neutrophil migration have shown both that cells can migrate up a concentration gradient of one chemoattractant in the presence of a saturating concentration of a second chemoattractant and that there are hierarchies of responses such that cells can migrate up the gradients of some chemoattractants even while moving down the gradients of others ^100, 101^. One or both processes could be operating in the sequential responses that we found to chemokines mediating arrest followed by TEM.

Finally, there is the question of how T cells that lack CCR2 might accomplish TEM *in vivo*. One prominent example of such trafficking is the migration of naïve T cells across HEVs into lymphoid organs, and considering this process is instructive. The principal receptor mediating T cell trafficking across HEVs is CCR7, which importantly, responds to its two ligands, CCL21 and CCL19, with differences in downstream signaling ^102^. Additionally, CCL21 contains a carboxy-terminal tail that binds avidly to GAGs, and CCL21 is produced by HEV endothelial cells, while CCL19 does not bind to GAGs and is produced not by HEV endothelial cells but by the surrounding stromal cells ^103, 104^. In studies of the roles for CCR7 and these ligands in the trafficking of dendritic cells within tissue, it was found that only the immobilized CCL21 could trigger integrin-mediated adhesion, whereas a soluble gradient of CCL19 (or a truncated CCL21) could induce the adherent cells to undergo directional migration ^105^. Although in this case one receptor would seem to be mediating both arrest and TEM on HEV endothelial cells, ligand-biased signaling would produce the functional equivalent of two receptors, one responding to a GAG-binding luminal ligand and the second responding to a soluble ligand produced on the abluminal side that we can reasonably presume forms a transendothelial concentration gradient. In this way both naïve cells and the multi-receptor, pro-inflammatory memory T cells that we have studied here can maintain the functional separation of arrest and TEM, mediated, respectively, by endothelial cell bound and non-bound chemokines. Depending on the endothelial cells, tissues, and biological responses, differences in chemokine production and GAG expression could enable analogous mechanisms to achieve T cell extravasation where receptors/ligands exchange roles, providing these cells with a broader repertoire of chemokine receptors to mediate TEM.

Although our work has been necessarily limited to *ex vivo* experiments, the particular strength of our studies is the integrated investigation of multiple aspects of the effector activities of bona fide human type 17 cells. We have identified a strong connection between the expression of CCR2 on type 17 cells and a pattern of gene and protein expression associated with pathogenicity in immune-mediated disease and host defense against mycobacteria - and we have shown that this same receptor is essential for migration across inflamed endothelium, a critical component of type 17 effector function. We have investigated in detail how multiple chemokine receptors and their ligands, expressed in complex and combinatorial patterns on human memory Th cells, are coordinated for extravasation. These studies have revealed fundamental principles of chemokine system functioning on lymphocytes, raising key questions for future research, and establishing a more informed basis for considering interventions to block or enhance T cell trafficking into tissue.

## Materials and methods

### Cell culture

Cryopreserved primary HUVECs were purchased from ATCC (PCS-100-013; ATCC). HUVECs were cultured in complete vascular cell growth medium (Vascular Cell Basal Medium: (PCS-100-030), supplemented with Endothelial Cell Growth Kit-VEGF: (PCS-100-041; ATCC) containing penicillin-streptomycin (100 U/ml, 15140122; Gibco) and maintained at 37°C in 5% CO2. To subculture, HUVECs were washed once with PBS, treated with 0.05% trypsin-EDTA (25300054; Thermo Fisher Scientific) for 90 s and were used until passage three. Human primary CD4^+^ T cells were cultured overnight in RPMI-10 media (RPMI 1640, 11875119; Gibco), penicillin-streptomycin (100 U/ml, 15140122; Gibco) and HEPES (10 mM, 15630080; Gibco) containing 10% FBS (100-500; Gemini Bio-product) at 37°C in 5% CO_2_.

### Purification of lymphocyte subsets from blood, cell sorting and culturing with cytokines

Human elutriated lymphocytes from healthy donors were obtained by the Department of Transfusion Medicine, Clinical Center, National Institutes of, under a protocol approved by the Institutional Review Board and after explanation of the risks and obtaining informed consent. Human CD4^+^ T cells were isolated from approximately 2 x 10^8^ elutriated lymphocytes to approximately 90% purity by negative selection using RosetteSep human CD4^+^ T cell enrichment cocktail (15062; Stem Cell Technologies), using Ficoll/Hypaque (45-001-749; Cytiva) density centrifugation. If needed, red blood cells were removed using ACK lysing buffer (BP10-548E; Lonza). Cells were incubated with anti-human CCR2-biotin (FAB151B; R&D Systems/ 357224; BioLegend) in FACS buffer (HBSS, 14025092; Gibco plus 4% FBS) for 30 min at room temperature (RT) and following washing, cells were stained with streptavidin-PE (405245), anti– human CD4-Brilliant Violet 421 (317434), anti–human CD45RO-Brilliant Violet 605 (304238), anti-human CD62L-PE-Cy5 (304808), anti-human CD25-APCCy7 (302614), HLADR-APCCy7 (307618) anti-human CD127 FITC (351314), all from BioLegend and anti-human CCR6 PECy7 (560620; BD Biosciences) for 30 min at RT. Cell subsets were sorted to nearly 100% purity using FACSAria II Cell Sorter (BD Biosciences). Purified cells from five subsets were cultured in IL-12 (20 ng/ml, 219-IL-005; R&D Systems), IL-18 (20 ng/ml 9124-IL-010; R&D Systems), IL-23 (20 ng/ml, 1290-IL-010/CF; R&D Systems), IL-1β (20 ng/ml 201-LB-005/CF; R&D Systems) and TNF-α (20 ng/ml 210-TA-005/CF; R&D Systems) in media containing 10 ng/ml IL-7 (BT-007-010; R&D Systems) at 37°C in 5% CO_2_ and after 3.5 days, supernatants were collected and concentrations of IFN-γ and GM-CSF were determined using ELISA kits from R&D Systems (IFN-γ; DIF50C, GM-CSF; DGM00) according to the manufacturer’s instructions.

### Analysis of CD4^+^ T cell migration under flow

μ-Slide I ^0.4^ Luer parallel plate flow chambers (80176; Ibidi) were coated with fibronectin (50 μg/ml, 1918-FN; R&D Systems) in PBS and HUVECs were plated at confluence and were stimulated with 40 ng/ml recombinant human TNF-α (210-TA; R&D Systems) and/or 50 ng/ml IFN-γ (285-IF; R&D Systems) for 18–20 h at 37°C in 5% CO2. HUVEC-coated parallel plate flow chambers were assembled with a two-pump system (Harvard Apparatus). T cells were re-suspended at 4×10^5^ cells/ml in RPMI-10 media. Perfusion of T cells into the flow chambers was performed at 37°C under a shear stress 0.75 dyn/cm^2^ for 4 min to allow accumulation of T cells, followed by a constant shear stress of 3 dyn/cm^2^ for 16 min. Images were acquired at a rate of four frames per second with an integrated fluorescence microscope, Leica AF 6000LX (Leica Microsystems Inc.) with a 20 x DIC objective. Data were analyzed using Imaris (Oxford Instruments) to track rolling, arrest and TEM. We categorized arresting cells as cells that remained stopped on the HUVEC monolayer for more than 10 s under a sheer stress of 3 dyn/cm^2^; rolling cells as cells that rolled before arresting (whether that arrest had initiated at 0.75 dyn/cm^2^ or at 3 dyne/cm^2^), and cells that rolled under a sheer stress of 3 dyn/cm^2^ but then detached; and transmigrating cells as cells that went underneath through stepwise darkening under a sheer stress of 3 dyn/cm^2^. For inhibiting Gi/o proteins, T cell were pre-incubated with 1 mg/ml pertussis toxin (3097; R&D Systems) in RPMI-10 medium for 3 h at 37°C. For blocking CCR2, cells were pre-incubated for 30 min with BMS CCR2 22 (2 μM, 3129), for CCR5, with Maraviroc (10 μM, 3756) and for blocking CXCR3 with AMG487 (10 μM, 4487) all from R&D Systems, at 37°C, and the inhibitors were left in the medium throughout the assays. For neutralizing the CCR6 ligand, CCL20, HUVEC monolayers in chambers slides were pre-treated with 20 mg/ml anti-human CCL20/MIP-3α antibody (MAB360; R&D Systems) for 2 h at 37°C, and antibody was maintained at 10 ng/ml throughout the assays. For treatment of T cells with CCL2, recombinant human CCL2 (100 ng/ml, 279-MC-010; R&D Systems) was added to the cells immediately before loading them into the flow chamber and the CCL2 concentration was maintained during the initial 4 min of the assay.

### Intracellular staining for flow cytometry

T cells purified by cell sorting were stimulated with Leukocyte Activation Cocktail, with GolgiPlus (550583; BD Pharmingen) for 6 h at 37°C with 5% CO_2_ before being stained with following antibodies alone or in combinations: anti-IL-17A (512305), anti-IFN-γ (502530), BioLegend), anti-TNF-α (502944), anti-GM-CSF (502305) all from BioLegend and anti-CCL20 (C360A; R&D Systems) by using Cytofix/CytoPerm Plus kit (555028; BD Pharmingen). Samples were analyzed using a Fortessa flow cytometer (BD Biosciences), and the data were analyzed using FlowJo software (BD Biosciences).

### Immunofluorescence and confocal microscopy

Circular cover slips (12 mm diameter) were placed in wells of a 24-well tissue culture plate and coated with recombinant human fibronectin for 2 h at 37°C or overnight at 4°C. Cells were grown on these cover slips for 18-20 h in complete vascular cell growth medium at 37^0^C with 5% CO_2_. Samples were fixed with paraformaldehyde (4%, Thermo Scientific)) in PBS and 2% sucrose for 30 min at RT. For permeabilization, samples were incubated with 0.1% Triton X-100 for 5 min and after extensive washing, blocked with 2% BSA/PBS for 30 min at RT. Samples were incubated with primary antibodies against chemokines CCL2 (5 μg, MAB679) CCL5 (5 μg, MAB278), CCL7 (5 μg, MAB282), CCL8 (5 μg, MAB281), CCL20 (5μg, AF360), CXCL9 (5 μg, MAB392), mouse IgG_1_ isotype control (5 μg, MAB002) and mouse IgG_2B_ isotype control (MAB004) all from R&D Systems, at RT for 2 h. Cells were washed three times with PBS before incubating with secondary antibody Alexa Fluor 488 (1:500, A-11017; Invitrogen) for 1 h and after washing three times with PBS, were counter-stained with DAPI (1μg/ml 62248; Invitrogen). Images were captured using a Leica SP8 (690) fluorescence microscope. Images were obtained with 40x objective. Images were analyzed using IMARIS software (Oxford Instruments).

### Chemokine binding assay

HUVECs were grown as non-stimulated or TNF-α-stimulated monolayer for 18 h on circular coverslips coated with fibronectin as described above. Chemically synthesized human CCL2 and CCL5 were supplied with site-specific biotinylation (Almac). 120 nM chemokine was added to the cells for 30 min at 37°C in 5% CO_2,_ after washing with Vascular Cell Basal Medium, cells were incubated with Alexa Fluor 594 streptavidin (1:500, 405240; BioLegend) for an additional 30 min. Cells were washed and fixed with 4% paraformaldehyde and after counter stained with DAPI images were captured using Leica SP8 (690) fluorescence microscope and analyzed using IMARIS software.

### Enzyme-Linked Immunosorbent Assay (ELISA)

For detection of chemokine secretion, HUVECs were left unstimulated or stimulated with TNF-α and/or IFN-γ for 18 h in culture conditions as described above. Medium was collected, and cells were lysed in medium containing 0.1% Triton X-100. Concentrations of chemokines were determined using ELISA kits from R&D Systems (CCL2; DCP00, CCL7; DCC700, CCL8; DY281, CCL20; DM3A00) according to the manufacturer’s instructions. Final values were averages of measurements performed in duplicate or triplicate. Levels of chemokines in the conditioned medium from cultured cells and the cell lysate were determined by reference to a standard curve produced using recombinant protein. To measure apically and basally secreted chemokines, HUVEC monolayers were grown overnight in culture conditions as described above but on fibronectin coated transwells (3415; Corning) with a 3 μm pore size. Monolayers were washed gently and activated with TNF-α and IFN-γ and conditioned medium was collected from upper and lower wells after 4 h. The upper well samples were diluted with medium to equalize sample volumes and chemokine concentrations were measured by ELISA.

### Total RNA isolation and real-time RT-PCR

For detection of chemokine gene expression in HUVECs, cells were cultured as described above, either unstimulated or stimulated with TNF-α and/or IFN-γ for 18h, and total RNA was isolated using RNeasy mini kit (Qiagen) according to the manufacturer’s instructions. Real-time RT-PCR was performed using 50 ng of RNA and the Platinum Quantitative RT-PCR ThermoScript One-Step System with FAM primers and probe sets from Applied Biosystems (*CCL2*; Hs00234140_m1, *CCL5*; Hs00982282_m1, *CCL7*; Hs00171147_m1, *CCL8*; Hs04187715_m1, *CCL20*; Hs00355476_m1, *CXCL9*; Hs00171065_m1, *CXCL10*; Hs00171042_m1, *CXCL11*; Hs00171138_m1). Values for chemokine mRNAs were normalized based on the values for *GAPDH* (Hs02786624_g1).

### RNA-Seq

T cells were cultured unstimulated or stimulated with cell stimulation cocktail (00-4970-03; Invitrogen) for 3 h at 37°C in 5% CO_2_. Extraction of total RNA was done from all samples using RNeasy mini kit (Qiagen) according to the manufacturer’s instructions. Oligo dT-based mRNA enrichment was done using NEBNext® Poly(A) mRNA Magnetic Isolation Module (E7490L) and library preparation was completed using NEBNext® Ultra™ II Directional RNA Library Prep Kit for Illumina® (E7760L), with 50 ng total RNA input per sample and targeting for 200 bp RNA insert size and NEBNext® Multiplex Oligos for Illumina (Unique Dual Index Primers) (E6440L). Sequencing was done using an Illumina NextSeq 500 system. The resulting FastQ files were checked for the quality control using FastQC (Babraham Bioinformatics tool). Adapter trimming was done using trimmomatic (v0.32, PMID:23104886); alignment to the human reference genome was done with STAR (PMID:35311944); duplicate reads were removed using Picard MarkDuplicates tool; and number of reads per gene per sample were counted using the the featureCounts software from the subread package (PMID:24227677). Differential gene expression analysis was done in R (v4.2.1) using DESeq2 package (v1.38.1) ^106^ and results were filtered for genes with <5% probability of being false positive (adjusted P value <0.05).

### Single-cell RNA sequencing (scRNA-seq)

CCR6^+^CCR2^+^CD4^+^ and total unselected memory T cells were stimulated with cell stimulation cocktail (00-4970-03; Invitrogen) for 3 h at 37°C in 5% CO_2_. Cells were loaded onto the Chromium Single Cell Library & Gel Bead Kit v3.1 (10x Genomics), to generate libraries for scRNA-seq and sequenced using the HiSeq 4000 System (Illumina). Each library was down sampled to equivalent sequencing depth and libraries were merged with cellranger aggr v2.0.2 pipeline (10X Genomics). Data were imported into R Studio (R 3.6.2), and the Seurat package (Seurat 3.1.4, https://github.com/satijalab/seurat) was used to process and analyze the gene expression data. After quality control, genes expressed in fewer than 10 cells were filtered out, and cells with more than 500 genes detected and fewer than 5% mitochondrial genes were retained for analysis. Data were standardized and normalized, and the principal component analysis (PCA) was implemented for nonlinear dimension reduction. Finally, cluster analysis was used to identify cell subtypes and UMAP was used for visualization of dimension reduction results. The FindAllMarkers function in the Seurat package was used to identify the significantly differentially expressed genes (DEGs) of every cluster vs. all others, and genes with the top 10 log2 fold-changes (log2FC) for every cluster were displayed as a heatmap. Monocle 2 (http://cole-trapnell-lab.github.io/monocle-release) was used to conduct the pseudo-time analysis in the cells. Monocle 2 used an algorithm to learn the changes in gene expression sequences that each cell must go through as part of a dynamic biological process (differentiation, for example). Cell dimensionality was reduced by the DDRTree method, and cells were sequenced in pseudotime and finally visualized.

### Transduction of HUVECs

HUVECs were grown overnight in culture conditions as described above with 0.2 x 10^6^ cells per well of a 6-well tissue culture plate, and the following day cells were washed and left in complete vascular cell growth medium containing 8 μg/ml polybrene for two hours before adding control and CXCL9-CCL2 chimera expressing lentivirus particles at a multiplicity of infection of five. Lentiviral particles expressing CCL2-CXCL9 fusion protein and negative control were purchased from GeneCopoeia.

### CCR2 internalization assay

PBMC were isolated from whole blood and treated with recombinant human CCL2 (279-MC-010; R&D Systems) at 100 ng/ml, 1 mg/ml and 10 mg/ml and cells were stained with anti-human CD4-APC, CD45RO-BV605, CCR2-PE and CXCR3-PE-Cy7 as described above, except that CCL2 was maintained at the indicated concentrations during the staining. Samples were analyzed using a Fortessa flow cytometer (BD Biosciences), and the data were analyzed using FlowJo software (BD Biosciences).

### Statistical analysis

Statistical analyses were performed using GraphPad Prism or R. Wilcoxon matched pairs signed rank test was used for the data from six or more experiments. Paired Student’s t tests were used for statistical analysis of the data from less than six and more than three experiments. Pooled experiments are represented as median or mean ± SEM, as indicated. The level of significance was depicted as * P < 0.05, ** P < 0.01, *** P < 0.001, and ns (not statistically significant). All experiments were performed ≥2 times, as indicated within the figure legend for each corresponding dataset.

## Acknowledgements

We are grateful to Tracy M. Handel, UC San Diego, for helpful advice, Jean K. Lim, Icahn School of Medicine at Mount Sinai, for critical reading of the manuscript and the members of the Research Technologies Branch, NIAID, for their help with cell sorting. This research was supported by the Intramural Research Program of the NIAID, NIH. The authors declare no competing financial interests.

## References

1. Devarajan, P. & Chen, Z. Autoimmune effector memory T cells: the bad and the good. Immunol Res 57, 12–22 (2013).

2. Remakus, S. & Sigal, L.J. Memory CD8(+) T cell protection. Adv Exp Med Biol 785, 77–86 (2013).

3. Sattler, A. et al. Cytokine-induced human IFN-gamma-secreting effector-memory Th cells in chronic autoimmune inflammation. Blood 113, 1948–1956 (2009).

4. Teijaro, J.R. et al. Cutting edge: Tissue-retentive lung memory CD4 T cells mediate optimal protection to respiratory virus infection. J Immunol 187, 5510–5514 (2011).

5. Teijaro, J.R., Verhoeven, D., Page, C.A., Turner, D. & Farber, D.L. Memory CD4 T cells direct protective responses to influenza virus in the lungs through helper-independent mechanisms. J Virol 84, 9217–9226 (2010).

6. Campbell, D.J., Kim, C.H. & Butcher, E.C. Separable effector T cell populations specialized for B cell help or tissue inflammation. Nat Immunol 2, 876–881 (2001).

7. Kaech, S.M., Wherry, E.J. & Ahmed, R. Effector and memory T-cell differentiation: implications for vaccine development. Nat Rev Immunol 2, 251–262 (2002).

8. Gebhardt, T. et al. Different patterns of peripheral migration by memory CD4+ and CD8+ T cells. Nature 477, 216–219 (2011).

9. Glennie, N.D., Volk, S.W. & Scott, P. Skin-resident CD4+ T cells protect against Leishmania major by recruiting and activating inflammatory monocytes. PLoS Pathog 13, e1006349 (2017).

10. Iijima, N. & Iwasaki, A. T cell memory. A local macrophage chemokine network sustains protective tissue-resident memory CD4 T cells. Science 346, 93–98 (2014).

11. Mora, J.R. & von Andrian, U.H. T-cell homing specificity and plasticity: new concepts and future challenges. Trends Immunol 27, 235–243 (2006).

12. Woodland, D.L. & Kohlmeier, J.E. Migration, maintenance and recall of memory T cells in peripheral tissues. Nat Rev Immunol 9, 153–161 (2009).

13. Zhu, J. T Helper Cell Differentiation, Heterogeneity, and Plasticity. Cold Spring Harb Perspect Biol 10 (2018).

14. Dong, C. Cytokine Regulation and Function in T Cells. Annu Rev Immunol 39, 51–76 (2021).

15. Lee, Y. et al. Induction and molecular signature of pathogenic TH17 cells. Nat Immunol 13, 991–999 (2012).

16. Stockinger, B. & Omenetti, S. The dichotomous nature of T helper 17 cells. Nat Rev Immunol 17, 535–544 (2017).

17. Singh, S.P., et al. Human CCR6+ Th cells show both an extended stable gradient of Th17 activity and imprinted plasticity. *bioRxiv*, 2023.2001.2005.522630 (2023).

18. Hedrick, M.N., Lonsdorf, A.S., Hwang, S.T. & Farber, J.M. CCR6 as a possible therapeutic target in psoriasis. Expert Opin Ther Targets 14, 911–922 (2010).

19. Reboldi, A. et al. C-C chemokine receptor 6-regulated entry of TH-17 cells into the CNS through the choroid plexus is required for the initiation of EAE. Nat Immunol 10, 514–523 (2009).

20. Baba, M. et al. Identification of CCR6, the specific receptor for a novel lymphocyte-directed CC chemokine LARC. J Biol Chem 272, 14893–14898 (1997).

21. Liao, F. et al. STRL22 is a receptor for the CC chemokine MIP-3alpha. Biochem Biophys Res Commun 236, 212–217 (1997).

22. Zhang, H.H. et al. CCR2 identifies a stable population of human effector memory CD4+ T cells equipped for rapid recall response. J Immunol 185, 6646–6663 (2010).

23. Hedrick, M.N. et al. CCR6 is required for IL-23-induced psoriasis-like inflammation in mice. J Clin Invest 119, 2317–2329 (2009).

24. Verma, A.H., et al. Oral epithelial cells orchestrate innate type 17 responses to Candida albicans through the virulence factor candidalysin. Sci Immunol 2 (2017).

25. Schall, T.J. & Proudfoot, A.E. Overcoming hurdles in developing successful drugs targeting chemokine receptors. Nat Rev Immunol 11, 355–363 (2011).

26. Springer, T.A. Traffic signals for lymphocyte recirculation and leukocyte emigration: the multistep paradigm. Cell 76, 301–314 (1994).

27. Lee, C.H. et al. C/EBPdelta drives interactions between human MAIT cells and endothelial cells that are important for extravasation. Elife 7 (2018).

28. Acosta-Rodriguez, E.V. et al. Surface phenotype and antigenic specificity of human interleukin 17-producing T helper memory cells. Nat Immunol 8, 639–646 (2007).

29. Singh, S.P., Zhang, H.H., Foley, J.F., Hedrick, M.N. & Farber, J.M. Human T cells that are able to produce IL-17 express the chemokine receptor CCR6. J Immunol 180, 214–221 (2008).

30. Ramesh, R. et al. Pro-inflammatory human Th17 cells selectively express P-glycoprotein and are refractory to glucocorticoids. J Exp Med 211, 89–104 (2014).

31. Karmaus, P.W.F. et al. Metabolic heterogeneity underlies reciprocal fates of TH17 cell stemness and plasticity. Nature 565, 101–105 (2019).

32. Schnell, A. et al. Stem-like intestinal Th17 cells give rise to pathogenic effector T cells during autoimmunity. Cell 184, 6281–6298 e6223 (2021).

33. Shulman, Z. et al. Transendothelial migration of lymphocytes mediated by intraendothelial vesicle stores rather than by extracellular chemokine depots. Nat Immunol 13, 67–76 (2011).

34. Park, M.K. et al. The CXC chemokine murine monokine induced by IFN-gamma (CXC chemokine ligand 9) is made by APCs, targets lymphocytes including activated B cells, and supports antibody responses to a bacterial pathogen in vivo. J Immunol 169, 1433–1443 (2002).

35. Vanheule, V. et al. The Positively Charged COOH-terminal Glycosaminoglycan-binding CXCL9(74-103) Peptide Inhibits CXCL8-induced Neutrophil Extravasation and Monosodium Urate Crystal-induced Gout in Mice. J Biol Chem 290, 21292–21304 (2015).

36. Harbour, S.N., Maynard, C.L., Zindl, C.L., Schoeb, T.R. & Weaver, C.T. Th17 cells give rise to Th1 cells that are required for the pathogenesis of colitis. Proc Natl Acad Sci U S A 112, 7061–7066 (2015).

37. Sallusto, F., Cassotta, A., Hoces, D., Foglierini, M. & Lanzavecchia, A. Do Memory CD4 T Cells Keep Their Cell-Type Programming: Plasticity versus Fate Commitment? T-Cell Heterogeneity, Plasticity, and Selection in Humans. Cold Spring Harb Perspect Biol 10 (2018).

38. Hirota, K. et al. Autoimmune Th17 Cells Induced Synovial Stromal and Innate Lymphoid Cell Secretion of the Cytokine GM-CSF to Initiate and Augment Autoimmune Arthritis. Immunity 48, 1220–1232 e1225 (2018).

39. Arlehamn, C.L. et al. Transcriptional profile of tuberculosis antigen-specific T cells reveals novel multifunctional features. J Immunol 193, 2931–2940 (2014).

40. Nathan, A. et al. Multimodally profiling memory T cells from a tuberculosis cohort identifies cell state associations with demographics, environment and disease. Nat. Immunol. 22, 781-+ (2021).

41. Strickland, N. et al. Characterization of Mycobacterium tuberculosis-Specific Cells Using MHC Class II Tetramers Reveals Phenotypic Differences Related to HIV Infection and Tuberculosis Disease. J. Immunol. 199, 2440–2450 (2017).

42. Okada, S., et al. IMMUNODEFICIENCIES. Impairment of immunity to Candida and Mycobacterium in humans with bi-allelic RORC mutations. Science 349, 606–613 (2015).

43. Hartmann, F.J. et al. Multiple sclerosis-associated IL2RA polymorphism controls GM-CSF production in human TH cells. Nat Commun 5, 5056 (2014).

44. Restorick, S.M. et al. CCR6(+) Th cells in the cerebrospinal fluid of persons with multiple sclerosis are dominated by pathogenic non-classic Th1 cells and GM-CSF-only-secreting Th cells. Brain Behav Immun 64, 71–79 (2017).

45. Cao, Y. et al. Functional inflammatory profiles distinguish myelin-reactive T cells from patients with multiple sclerosis. Sci Transl Med 7, 287ra274 (2015).

46. Paulissen, S.M., van Hamburg, J.P., Dankers, W. & Lubberts, E. The role and modulation of CCR6+ Th17 cell populations in rheumatoid arthritis. Cytokine 74, 43–53 (2015).

47. McGeachy, M.J. et al. The interleukin 23 receptor is essential for the terminal differentiation of interleukin 17-producing effector T helper cells in vivo. Nat Immunol 10, 314–324 (2009).

48. El-Behi, M. et al. The encephalitogenicity of T(H)17 cells is dependent on IL-1- and IL- 23-induced production of the cytokine GM-CSF. Nat Immunol 12, 568–575 (2011).

49. Hasan, Z. et al. JunB is essential for IL-23-dependent pathogenicity of Th17 cells. Nat Commun 8, 15628 (2017).

50. Lin, C.C. et al. IL-1-induced Bhlhe40 identifies pathogenic T helper cells in a model of autoimmune neuroinflammation. J Exp Med 213, 251–271 (2016).

51. Mazzoni, A. et al. Eomes controls the development of Th17-derived (non-classic) Th1 cells during chronic inflammation. Eur J Immunol 49, 79–95 (2019).

52. Rasouli, J., et al. A distinct GM-CSF(+) T helper cell subset requires T-bet to adopt a TH1 phenotype and promote neuroinflammation. Sci Immunol 5 (2020).

53. Emming, S. et al. A molecular network regulating the proinflammatory phenotype of human memory T lymphocytes. Nat Immunol 21, 388–399 (2020).

54. Komuczki, J. et al. Fate-Mapping of GM-CSF Expression Identifies a Discrete Subset of Inflammation-Driving T Helper Cells Regulated by Cytokines IL-23 and IL-1beta. Immunity 50, 1289–1304 e1286 (2019).

55. Pierson, E.R. & Goverman, J.M. GM-CSF is not essential for experimental autoimmune encephalomyelitis but promotes brain-targeted disease. JCI Insight 2, e92362 (2017).

56. Kara, E.E. et al. CCR2 defines in vivo development and homing of IL-23-driven GM-CSF-producing Th17 cells. Nat Commun 6, 8644 (2015).

57. Croxford, A.L. et al. The Cytokine GM-CSF Drives the Inflammatory Signature of CCR2+ Monocytes and Licenses Autoimmunity. Immunity 43, 502–514 (2015).

58. Codarri, L. et al. RORgammat drives production of the cytokine GM-CSF in helper T cells, which is essential for the effector phase of autoimmune neuroinflammation. Nat Immunol 12, 560–567 (2011).

59. Hou, L. & Yuki, K. CCR6 and CXCR6 Identify the Th17 Cells With Cytotoxicity in Experimental Autoimmune Encephalomyelitis. Front Immunol 13, 819224 (2022).

60. Lee, H.-G. et al. Pathogenic function of bystander-activated memory-like CD4+ T cells in autoimmune encephalomyelitis. Nature Communications 10, 709 (2019).

61. Izikson, L., Klein, R.S., Charo, I.F., Weiner, H.L. & Luster, A.D. Resistance to experimental autoimmune encephalomyelitis in mice lacking the CC chemokine receptor (CCR)2. J Exp Med 192, 1075–1080 (2000).

62. Fife, B.T., Huffnagle, G.B., Kuziel, W.A. & Karpus, W.J. CC chemokine receptor 2 is critical for induction of experimental autoimmune encephalomyelitis. J Exp Med 192, 899–905 (2000).

63. Sato, W. et al. CCR2(+)CCR5(+) T cells produce matrix metalloproteinase-9 and osteopontin in the pathogenesis of multiple sclerosis. J Immunol 189, 5057–5065 (2012).

64. Wilson, N.J. et al. Development, cytokine profile and function of human interleukin 17-producing helper T cells. Nat Immunol 8, 950–957 (2007).

65. Lee, W.W. et al. Regulating human Th17 cells via differential expression of IL-1 receptor. Blood 115, 530–540 (2010).

66. Duhen, T. & Campbell, D.J. IL-1beta promotes the differentiation of polyfunctional human CCR6+CXCR3+ Th1/17 cells that are specific for pathogenic and commensal microbes. J Immunol 193, 120–129 (2014).

67. Dorner, B.G. et al. MIP-1alpha, MIP-1beta, RANTES, and ATAC/lymphotactin function together with IFN-gamma as type 1 cytokines. Proc Natl Acad Sci U S A 99, 6181–6186 (2002).

68. Rogge, L. et al. Transcript imaging of the development of human T helper cells using oligonucleotide arrays. Nat Genet 25, 96–101 (2000).

69. Schrum, S., Probst, P., Fleischer, B. & Zipfel, P.F. Synthesis of the CC-chemokines MIP-1alpha, MIP-1beta, and RANTES is associated with a type 1 immune response. J Immunol 157, 3598–3604 (1996).

70. Kebir, H. et al. Human TH17 lymphocytes promote blood-brain barrier disruption and central nervous system inflammation. Nat Med 13, 1173–1175 (2007).

71. Agak, G.W. et al. Extracellular traps released by antimicrobial T(H)17 cells contribute to host defense. Journal of Clinical Investigation 131 (2021).

72. Raveney, B.J. et al. Eomesodermin-expressing T-helper cells are essential for chronic neuroinflammation. Nat Commun 6, 8437 (2015).

73. Park, S., Anderson, N.L., Canaria, D.A. & Olson, M.R. Granzyme-Producing CD4 T Cells in Cancer and Autoimmune Disease. Immunohorizons 5, 909–917 (2021).

74. Chemin, K. et al. EOMES-positive CD4(+) T cells are increased in PTPN22 (1858T) risk allele carriers. Eur J Immunol 48, 655–669 (2018).

75. Masopust, D. & Picker, L.J. Hidden memories: frontline memory T cells and early pathogen interception. J Immunol 188, 5811–5817 (2012).

76. Barzilai, S. et al. M-sec regulates polarized secretion of inflammatory endothelial chemokines and facilitates CCL2-mediated lymphocyte transendothelial migration. J Leukoc Biol 99, 1045–1055 (2016).

77. Shulman, Z. et al. Lymphocyte crawling and transendothelial migration require chemokine triggering of high-affinity LFA-1 integrin. Immunity 30, 384–396 (2009).

78. Oynebraten, I. et al. Oligomerized, filamentous surface presentation of RANTES/CCL5 on vascular endothelial cells. Sci Rep 5, 9261 (2015).

79. Monneau, Y., Arenzana-Seisdedos, F. & Lortat-Jacob, H. The sweet spot: how GAGs help chemokines guide migrating cells. J Leukoc Biol 99, 935–953 (2016).

80. Bao, X. et al. Endothelial heparan sulfate controls chemokine presentation in recruitment of lymphocytes and dendritic cells to lymph nodes. Immunity 33, 817–829 (2010).

81. Tsuboi, K. et al. Role of high endothelial venule-expressed heparan sulfate in chemokine presentation and lymphocyte homing. J Immunol 191, 448–455 (2013).

82. Wang, L., Fuster, M., Sriramarao, P. & Esko, J.D. Endothelial heparan sulfate deficiency impairs L-selectin-and chemokine-mediated neutrophil trafficking during inflammatory responses. Nat Immunol 6, 902–910 (2005).

83. Kuschert, G.S. et al. Glycosaminoglycans interact selectively with chemokines and modulate receptor binding and cellular responses. Biochemistry 38, 12959–12968 (1999).

84. Nonaka, M. et al. Synthetic di-sulfated iduronic acid attenuates asthmatic response by blocking T-cell recruitment to inflammatory sites. Proc Natl Acad Sci U S A 111, 8173–8178 (2014).

85. Schenauer, M.R., Yu, Y., Sweeney, M.D. & Leary, J.A. CCR2 chemokines bind selectively to acetylated heparan sulfate octasaccharides. J Biol Chem 282, 25182–25188 (2007).

86. Lau, E.K. et al. Identification of the glycosaminoglycan binding site of the CC chemokine, MCP-1: implications for structure and function in vivo. J Biol Chem 279, 22294–22305 (2004).

87. Salanga, C.L. et al. Multiple glycosaminoglycan-binding epitopes of monocyte chemoattractant protein-3/CCL7 enable it to function as a non-oligomerizing chemokine. J Biol Chem 289, 14896–14912 (2014).

88. Randolph, G.J. & Furie, M.B. A soluble gradient of endogenous monocyte chemoattractant protein-1 promotes the transendothelial migration of monocytes in vitro. J Immunol 155, 3610–3618 (1995).

89. Weber, K.S., von Hundelshausen, P., Clark-Lewis, I., Weber, P.C. & Weber, C. Differential immobilization and hierarchical involvement of chemokines in monocyte arrest and transmigration on inflamed endothelium in shear flow. Eur J Immunol 29, 700–712 (1999).

90. Wagner, L. et al. Beta-chemokines are released from HIV-1-specific cytolytic T-cell granules complexed to proteoglycans. Nature 391, 908–911 (1998).

91. Palframan, R.T. et al. Inflammatory chemokine transport and presentation in HEV: a remote control mechanism for monocyte recruitment to lymph nodes in inflamed tissues. J Exp Med 194, 1361–1373 (2001).

92. Stoler-Barak, L., Barzilai, S., Zauberman, A. & Alon, R. Transendothelial migration of effector T cells across inflamed endothelial barriers does not require heparan sulfate proteoglycans. Int Immunol 26, 315–324 (2014).

93. Celie, J.W. et al. Subendothelial heparan sulfate proteoglycans become major L-selectin and monocyte chemoattractant protein-1 ligands upon renal ischemia/reperfusion. Am J Pathol 170, 1865–1878 (2007).

94. Girbl, T. et al. Distinct Compartmentalization of the Chemokines CXCL1 and CXCL2 and the Atypical Receptor ACKR1 Determine Discrete Stages of Neutrophil Diapedesis. Immunity 49, 1062–1076 e1066 (2018).

95. Carr, M.W., Roth, S.J., Luther, E., Rose, S.S. & Springer, T.A. Monocyte chemoattractant protein 1 acts as a T-lymphocyte chemoattractant. Proc Natl Acad Sci U S A 91, 3652–3656 (1994).

96. Miyabe, Y., Miyabe, C., Mani, V., Mempel, T.R. & Luster, A.D. Atypical complement receptor C5aR2 transports C5a to initiate neutrophil adhesion and inflammation. Sci Immunol 4 (2019).

97. Schwartz, A.B. et al. Elucidating the Biomechanics of Leukocyte Transendothelial Migration by Quantitative Imaging. Front. Cell. Dev. Biol. 9, 15 (2021).

98. Gardner, L., Patterson, A.M., Ashton, B.A., Stone, M.A. & Middleton, J. The human Duffy antigen binds selected inflammatory but not homeostatic chemokines. Biochem Biophys Res Commun 321, 306–312 (2004).

99. Yeh, Y.T. et al. Three-dimensional forces exerted by leukocytes and vascular endothelial cells dynamically facilitate diapedesis. Proc Natl Acad Sci U S A 115, 133–138 (2018).

100. Foxman, E.F., Kunkel, E.J. & Butcher, E.C. Integrating conflicting chemotactic signals. The role of memory in leukocyte navigation. J Cell Biol 147, 577–588 (1999).

101. Foxman, E.F., Campbell, J.J. & Butcher, E.C. Multistep navigation and the combinatorial control of leukocyte chemotaxis. J Cell Biol 139, 1349–1360 (1997).

102. Kohout, T.A. et al. Differential desensitization, receptor phosphorylation, beta-arrestin recruitment, and ERK1/2 activation by the two endogenous ligands for the CC chemokine receptor 7. J Biol Chem 279, 23214–23222 (2004).

103. de Paz, J.L. et al. Profiling heparin-chemokine interactions using synthetic tools. ACS Chem Biol 2, 735–744 (2007).

104. Luther, S.A., Tang, H.L., Hyman, P.L., Farr, A.G. & Cyster, J.G. Coexpression of the chemokines ELC and SLC by T zone stromal cells and deletion of the ELC gene in the plt/plt mouse. Proc Natl Acad Sci U S A 97, 12694–12699 (2000).

105. Schumann, K. et al. Immobilized chemokine fields and soluble chemokine gradients cooperatively shape migration patterns of dendritic cells. Immunity 32, 703–713 (2010).

106. Love, M.I., Huber, W. & Anders, S. Moderated estimation of fold change and dispersion for RNA-seq data with DESeq2. Genome Biol 15, 550 (2014).

