## Supplemental Figures for "Chemokine positioning determines mutually exclusive roles for their receptors in extravasation of pathogenic human T cells"

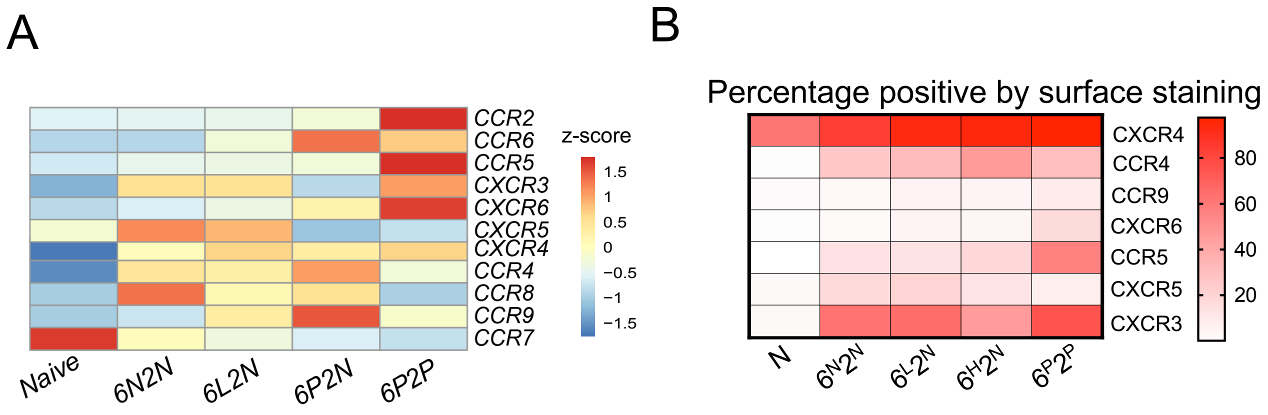


**Figure S1. Expression of additional chemokine receptors on the CD4$^+$ T cell subsets. (A)** Heatmap showing the expression levels of chemokine receptor mRNAs in cell subsets obtained from bulk RNA-seq from non-activated samples and in (B), averages of percentages of cells in the naïve and four memory subsets that were positive by surface staining for the indicated chemokine receptors from three donors. Staining with an isotype-matched control antibody was used as a negative control to determine the percentage of cells staining positive for a chemokine receptor.

**Figure S1**

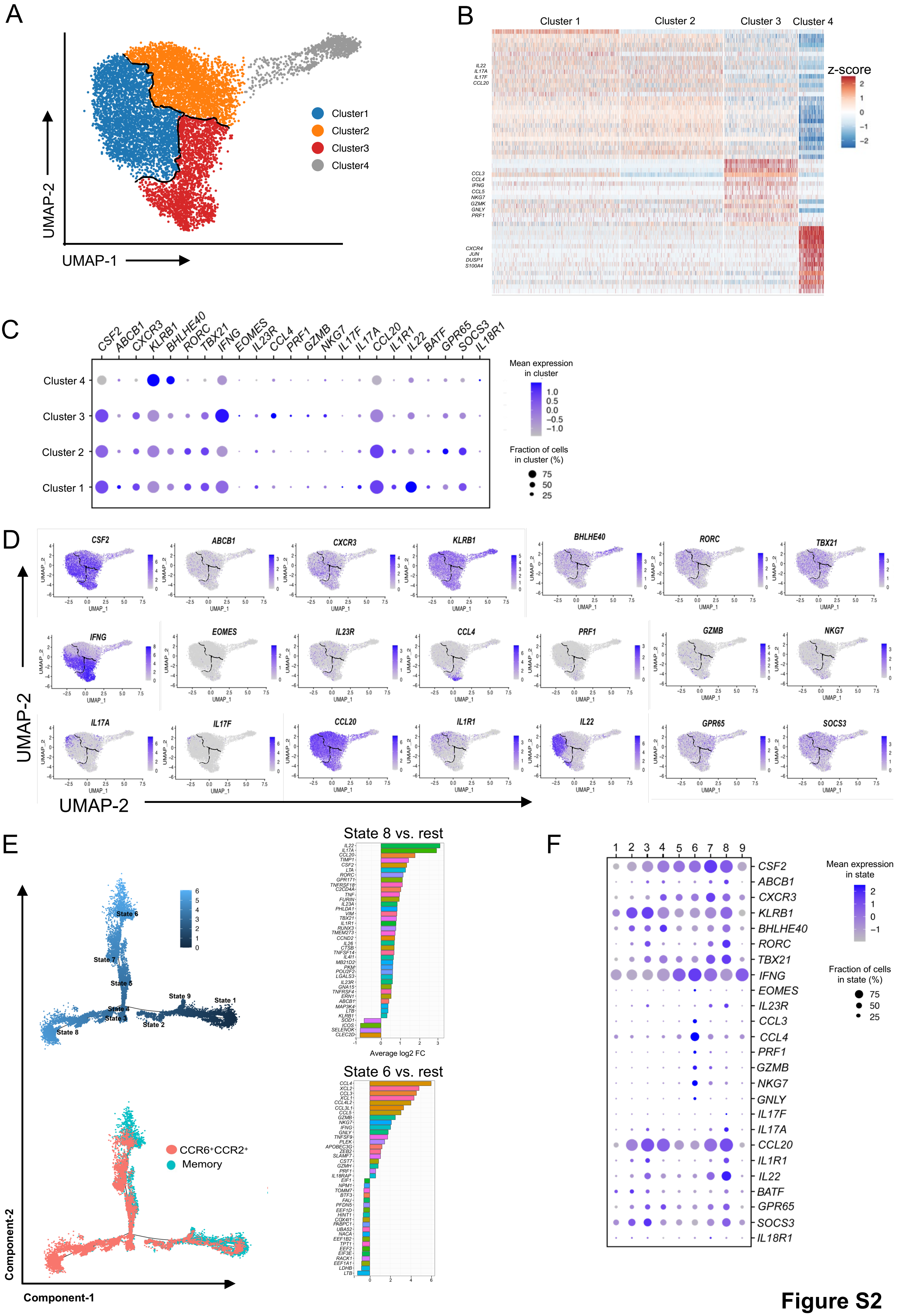
A
UMAP-2
UMAP-1
Cluster1
Cluster2
Cluster3
Cluster4
B
Cluster 1
Cluster 2
Cluster 3
Cluster 4
z-score
C
Mean expression in cluster
Fraction of cells in cluster (%)
Cluster 4
Cluster 3
Cluster 2
Cluster 1
D
UMAP-2
E
State 8 vs. rest
Average log2 FC
State 6 vs. rest
CCR6+CCR2+
Memory
Component-1
Component-2
F
Mean expression in state
Fraction of cells in state (%)

**Figure S2**

**Figure S2. CCR6$^+$CCR2$^+$CD4$^+$ T cells separate into clusters expressing *IL17A/F* vs. *IFNG*.** (A) UMAP projection and clustering from scRNA-seq profiles of 8,907 CCR6$^+$CCR2$^+$ cells from Donor 2 where each dot represents a single cell, with color codes of clusters defining cells with similar transcriptional profiles. (B) Heatmap indicating the upregulated and downregulated genes in each cluster compared to all other clusters. Rows represent genes and columns represent cells grouped by cluster, and some marker genes for clusters 1, 3 and 4 are listed on the left. Color coding reflects standardized gene expression values (z-scores). (C). Dot plot displaying gene expression for a subset of the genes displayed in (B). (D) Feature plots of expression levels of selected genes in cells in the UMAP scatter plot. Cluster borders are marked in black. (E) Pseudotime trajectories of single cell transcriptomic data showing states (upper left ) and with cells colored according to their origin as either total unselected memory or CCR6$^+$CCR2$^+$ cells (bottom left ). Right panel shows average log2 fold changes of genes with greatest differential expression in state 8 (top right) or state 6 (bottom right) versus the remaining cells. (F) Dot plot shows the expression of selected genes in each state.

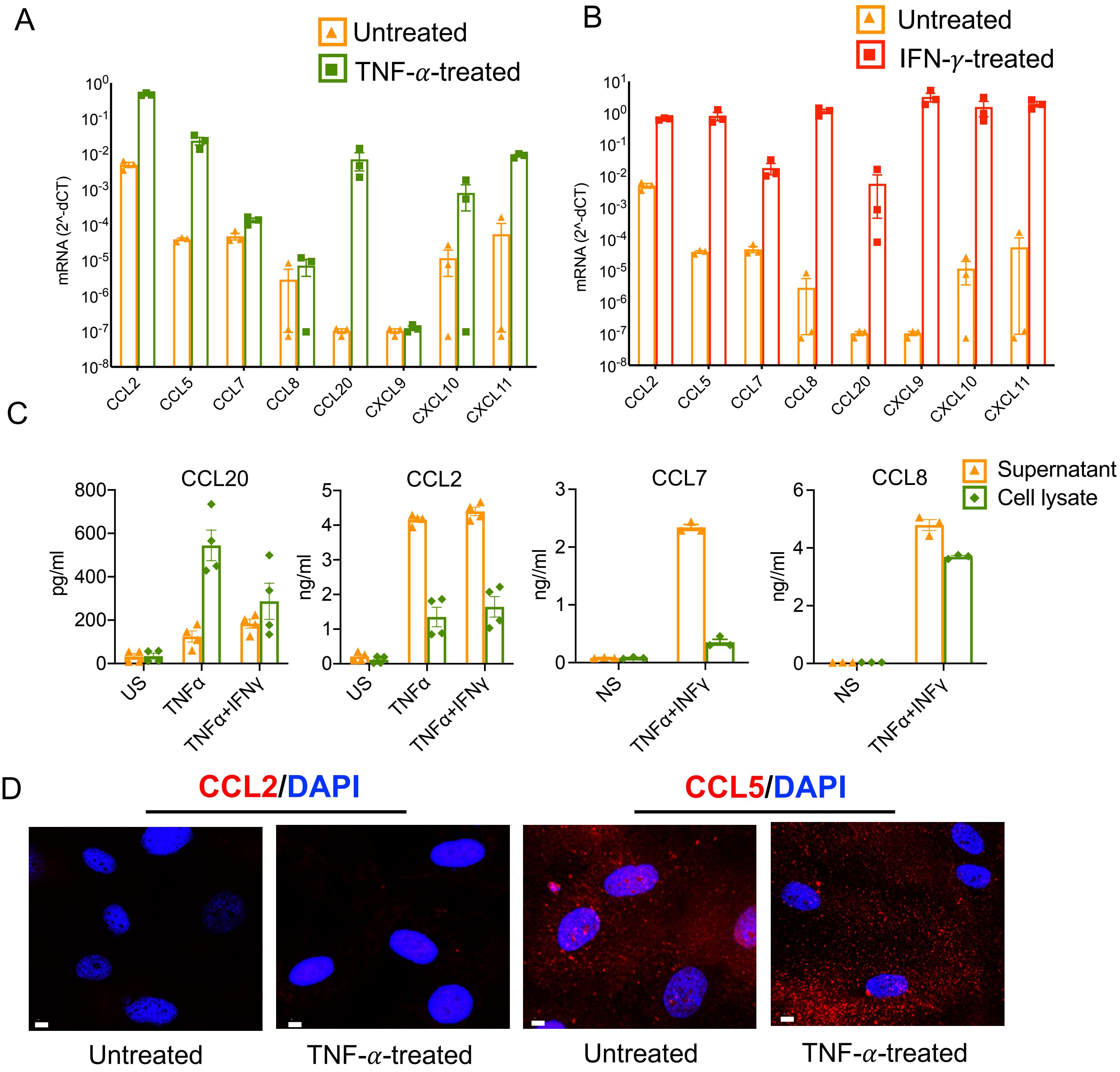


**Figure S3. CCR2 ligands produced by activated HUVECs are less cell-associated.** (A and B) Real-time RT-PCR analysis of mRNA levels of chemokines in TNF-$\alpha$ or with TNF-$\alpha$ + IFN-$\gamma$-treated HUVECs. (C) Culture supernatants collected from the TNF-$\alpha$ or TNF-$\alpha$ + IFN-$\gamma$-treated HUVECs were assayed for CCL20, CCL2, CCL7 and CCL8 by ELISA. Data are from at least three experiments and bars indicate means +/- SEM. (D) Confocal microscopy images at x40 magnification of non-permeabilized, untreated or TNF-$\alpha$-treated HUVECs after incubating with synthesized, CCL2 (left) or CCL5 (right) and visualizing using Alexa Fluor 594 streptavidin (red) as described in Materials and methods. The scale bars indicate 5 μm. Images are representative of at least three experiments.

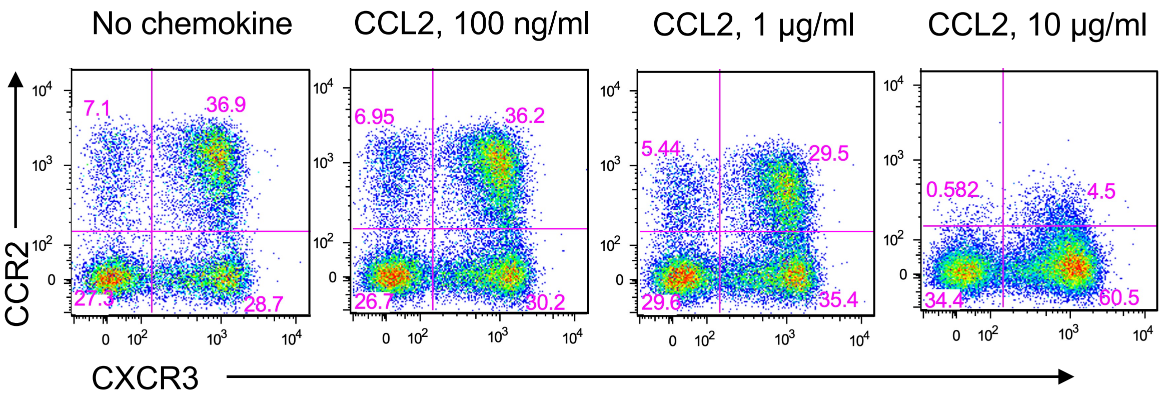


**Figure S4. CCR2 is not internalized on CD4⁺ T cells by 100 ng/ml CCL2.** Human PBMC were treated with the indicated concentrations of CCL2 as described in Materials and methods and dot plots show staining for CCR2 and CXCR3 on $CD4^+CD45R0^+$ cells. Results are representative three experiments.

**Figure S4**
